# Genomic analysis of a parasite invasion: colonization of the Americas by the blood fluke, *Schistosoma mansoni*

**DOI:** 10.1101/2021.10.25.465783

**Authors:** Roy N. Platt, Winka Le Clec’h, Frédéric D. Chevalier, Marina McDew-White, Philip T. LoVerde, Rafael R. de Assis, Guilherme Oliveira, Safari Kinunghi, Amadou Garba Djirmay, Michelle L. Steinauer, Anouk Gouvras, Muriel Rabone, Fiona Allan, Bonnie L. Webster, Joanne P. Webster, Aidan Emery, David Rollinson, Timothy J. C. Anderson

## Abstract

*Schistosoma mansoni,* a snail-vectored, blood fluke that infects humans, was introduced into the Americas from Africa during the Trans-Atlantic slave trade. As this parasite shows strong specificity to the snail intermediate host, we expected that adaptation to S. American *Biomphalaria* spp. snails would result in population bottlenecks and strong signatures of selection. We scored 475,081 single nucleotide variants (SNVs) in 143 *S. mansoni* from the Americas (Brazil, Guadeloupe, and Puerto Rico) and Africa (Cameroon, Niger, Senegal, Tanzania, and Uganda), and used these data to ask: (i) Was there a population bottleneck during colonization? (ii) Can we identify signatures of selection associated with colonization? And (iii) what were the source populations for colonizing parasites? We found a 2.4-2.9-fold reduction in diversity and much slower decay in linkage disequilibrium (LD) in parasites from East to West Africa. However, we observed similar nuclear diversity and LD in West Africa and Brazil, suggesting no strong bottlenecks and limited barriers to colonization. We identified five genome regions showing selection in the Americas, compared with three in West Africa and none in East Africa, which we speculate may reflect adaptation during colonization. Finally, we infer that unsampled African populations from central African regions between Benin and Angola, with contributions from Niger, are likely the major source(s) for Brazilian *S. mansoni*. The absence of a bottleneck suggests that this is a rare case of a serendipitous invasion, where *S. mansoni* parasites were preadapted to the Americas and were able to establish with relative ease.

## Introduction

Genomic characterization of parasites and pathogens is increasingly used as an aid to traditional epidemiological methods in reconstructing transmission patterns (de Oliveira et al., 2020; Nadeau, Vaughan, Scire, Huisman, & Stadler, 2021). On a longer time-scale genomic data can be used to understand biological invasions of pathogens into new continents, just as these methods are used for investigating biological invasions in free-living organisms (Rius, Bourne, Hornsby, & Chapman, 2015; Sherpa & Després, 2021). Such methods can determine the colonization route, source population, number of colonization events, whether diversity is reduced during colonization and evidence for adaptation in colonizing populations. Examining the consequences of historic invasions can inform our understanding of extant invasions.

The Trans-Atlantic slave trade lasted from 1502-1888 when the last remaining slave ports in Brazil were shutdown (Bergad, 2007). During this time, more than 12 million people were trafficked from Africa to slave ports in the Americas, representing one of the largest forced migration events in human history (Eltis, 2001). Along with the human cargo, a number of human pathogens were introduced into the Americas as well. For example, Parvo and hepatitis B viruses were successfully introduced into the Americas and rapidly spread through native populations leading to large scale outbreaks (Guzmán-Solís et al., 2021). *Trypansoma brucei*, the etiological agent for African Sleeping Sickness, was also introduced but failed to establish due to the absence of its’ intermediate host; the Tsetse fly (Steverding, 2020). Today viable populations of pathogens including Herpes simplex virus 2 (Forni et al., 2020), yellow fever virus (Bryant, Holmes, & Barrett, 2007), the parasitic nematode *Wuchereria bancrofti* (Small et al., 2019), among others (Steverding, 2020) are all a direct result of introductions during the Trans-Atlantic slave trade. In some cases, the genetic signatures of the introduction are still visible. For example, genetic diversity in South American *Leishmania chagasi* populations is halved and the effective population size (*Ne*) is reduced from 43.6M to 15.5K when compared to source populations in Africa (Leblois, Kuhls, François, Schönian, & Wirth, 2011; Schwabl et al., 2021). Here, we focus on successful invasion of the human-parasitic trematode, *Schistosoma mansoni*.

*S. mansoni* is distributed from Oman, through sub-Saharan Africa, to the Caribbean and countries along the eastern coast of South America. Phylogenetic evidence indicates that *S. mansoni* in West Africa and the Americas are closely related (Crellen et al., 2016; Desprès, Imbert-Establet, & Monnerot, 1993; Fletcher, LoVerde, & Woodruff, 1981; Morgan et al., 2005; Webster et al., 2013) and these observations, along with demographic reconstructions (Crellen et al., 2016), indicate a recent origin of *S. mansoni* in the Americas. As a result, there is strong evidence that *S*. *mansoni* co-migrated to the Americas during the forced human migrations of the Trans-Atlantic slave trade (Files, 1951). Furthermore, reduced diversity in mitochondrial haplotypes (Desprès et al., 1993; Fletcher et al., 1981; Morgan et al., 2005; Webster et al., 2013) in South American *S. mansoni* suggests the presence of a bottleneck during parasite establishment.

Our central goal was to use parasite genomic data to investigate this human-mediated, biological invasion and the impacts of a relatively recent, trans-continental, migration event. Parasites in the genus *Schistosoma* have a complex life cycle involving human definitive hosts and snail intermediate hosts (reviewed in Anderson & Enabulele, 2021). Eggs are expelled in human feces (*S. man*soni and *S. japonicum*) or urine (*S. haematobium*). Larvae (miracidia) hatch in fresh water and infect receptive snails. Once inside the snail host, the schistosomes reproduce asexually, and 2^nd^ stage larvae (cercariae) are released back into the water where they infect humans, mature into adult worms, and restart their life cycle. *S. mansoni* is diploid, with a well characterized 363Mb genome (Berriman et al., 2009; International Helminth Genomes Consortium, 2019; Protasio et al., 2012), ZW sex determination, obligate sexual reproduction of adult worms, and a relatively long life-span; from 5-10 years (Fulford, Butterworth, Ouma, & Sturrock, 1995).

The distribution of the intermediate snail host is a major driver of schistosome distribution. *S. haematobium*, a sister-taxon to *S. mansoni*, infects a different snail host; *Bulinus* spp. While people infected with both *S. mansoni* and *S. haematobium* were transported to S. America, only *S. mansoni* would have been able to establish due to the presence of the *Biomphalaria* spp. intermediate snail host(s) prior to their arrival (Morgan et al., 2005). *S. mansoni* shows strong specificity for species, and even strains of snails in the genus *Biomphalaria* (Webster & Woolhouse, 1998), however the *Biomphalaria* species assemblages differ between the Americas and Africa (DeJong et al., 2001). *B. pfeifferi*, *B. sudanica*, and *B. alexandrina*, are the primary intermediate hosts in Africa (DeJong et al., 2001), while *glabrata*, *B. tenagophila*, *B. straminea*, are the known snail hosts in South America (Vidigal et al., 2000). *S. mansoni* infections can impact the reproductive viability of their snail hosts, and there are strong co-evolutionary interactions driving resistance to infection in snails and for infectivity in parasites (Davies, Webster, & Woolhous, 2001; Theron, Rognon, Gourbal, & Mitta, 2014; Webster, Gower, & Blair, 2004). Several schistsome resistance genes have been localized within the snail genome (Tennessen et al., 2020; Tennessen et al., 2015) and polymorphic loci in both snail and parasites are thought to determine compatibility between snail and parasite (Mitta et al., 2017; Webster & Woolhouse, 1998; Woolhouse & Webster, 2000). Based on these observations, we hypothesize that the adaptation to novel, *Biomphalaria* spp. hosts would place strong selective pressures on *S. mansoni* as it became established in the Americas.

Adult schistosomes live in the blood vessels, making them difficult to sample. Genome and exome sequencing of schistosomes is now possible using whole genome amplification of miracidia larvae isolated from feces or urine (Doyle et al., 2019; Le Clec’h et al., 2018; Shortt et al., 2017) and several genome scale population analyses have recently been published (Berger et al., 2021; Platt et al., 2019; Shortt et al., 2017). Our goal is to address the following questions with the available sequence data from both the Africa (Niger, Senegal, Uganda, and Tanzania) and the Americas (Caribbean, Brazil): (i) Are the genomic data consistent with a West African origin of colonizing schistosome populations?; (ii) is there evidence for genetic bottlenecks during colonization; (iii) are there genomic signatures suggesting adaptation of colonizing parasites to the Americas; (iv) can we determine the source country or countries for Americas parasite populations?

## Materials and Methods

### Data and sample information

We examined published exomic and genomic data from 178 individual *Schistosoma* samples/isolates, from multiple geographical locations, available from three studies (Berriman et al., 2009; Chevalier et al., 2019; Crellen et al., 2016). All exome data is from Chevalier *et al*. (2016) and Chevalier *et al*. (2019). These data were generated from individual larval miracidia hatched from *S. mansoni* eggs and preserved on FTA cards. Exome libraries were generated via whole genome amplification followed by targeted capture of the exome (Le Clec’h et al., 2018). This method specifically targets 95% (14.81 Mb) of the exome with 2x tiled probes. The whole genome sequence data came from adult worms cultured through laboratory rodents and snails for two or more generations before whole genome library prep and sequencing (Berriman et al., 2009; Crellen et al., 2016; International Helminth Genomes Consortium, 2019). Sample origins are shown in Figure 2. Detailed metadata is available for each sample in Supplemental Table 1 including country of origin, species identification, NCBI Short Read Archive (SRA) accession, *etc*.

**Figure 2.**
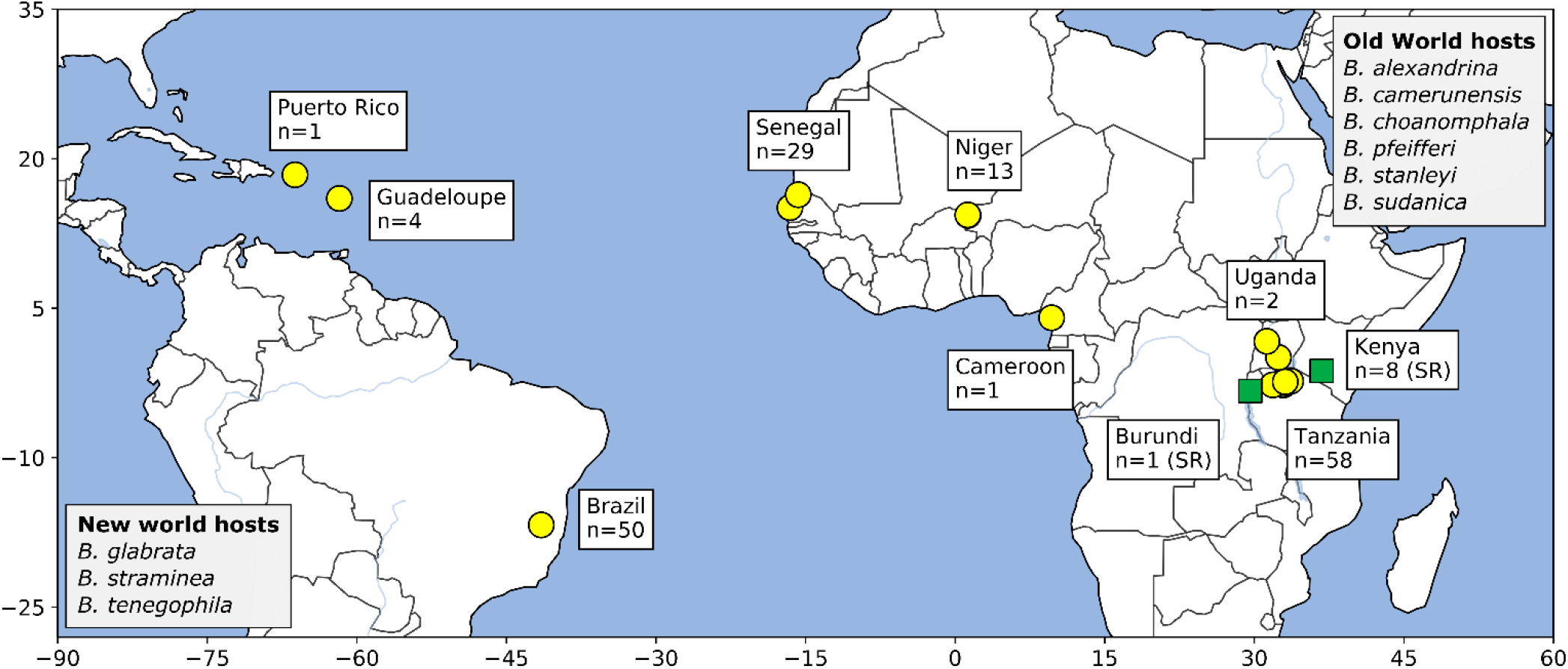
Sampling localities. Location and number of samples for *S. mansoni* (yellow circles) and *S. rodhaini* (green squares) samples included in this study. Members of the genus *Biomphalaria* are the predominant intermediate hosts with specific distributions of the different species involved in transmission varying across Africa and S. America. Intermediate snail vectors are listed in greyed boxes.

### Computational environment

We used conda v4.8.3 to manage virtual environments for all analyses. Sequence read filtering through genotyping steps were documented in a Snakemake v5.18.1 (Köster & Rahmann, 2012) workflow and all other analyses were performed in a series of Jupyter v1.0.0 notebooks. The code for this project including shell scripts, Snakemake workflows, notebooks, and environmental yaml files are available at https://github.com/nealplatt/sch_man_nwinvasion/releases/tag/v0.2 (last accessed 21 Oct 2021) and accessioned at https://doi.org/10.5281/zenodo.5590460 (last accessed 21 Oct 2021).

### Genotyping

Paired end reads were quality filtered with trimmomatic v0.39 (Bolger, Lohse, & Usadel, 2014) with the following parameters ‘LEADING:10 TRAILING:10 SLIDINGWINDOW:4:15 MINLEN:36’. Filtered reads were mapped to the *S. mansoni* genome (GenBank Assembly accession: GCA_000237925.3) using bwa v0.7.17-r1188 (Li & Durbin, 2010). We allowed up to 15 mismatches per 100 bp read (-n 15) to account for divergence between *S. mansoni* and *S. rodhaini*. Mapped single and paired reads were merged into a single file and all optical/PCR duplicates were removed with GATK v4.1.2.0’s (McKenna et al., 2010) MarkDuplicates. Single nucleotide variants (SNVs)were called with HaplotypeCaller and GenotypeGVCFs on a contig-by-contig basis, combined into a gvcf per individual, and finally merged into a single gvcf for the entire dataset. We used a high quality SNV data from Le Clec’h *et al*. (2021) as a training dataset for variant recalibration and scored SNV quality using ‘-an SOR -an MQ -an MQRankSum -an ReadPosRankSum’. Sensitivity Tranches (--truth-sensitivity-tranche) were set at 100, 99.5, 99, 97.5, 95, and 90. We re-calibrated SNVs using the 97.5 sensitivity tranche and filtered low confidence sites with the following set of filters (--filter-expression) QD < 2.0, MQ < 30.0, FS > 60.0, SOR > 3.0, MQRankSum < -12.5, and ReadPosRankSum < -8.0". All genotyping steps from read filtering through variant re-calibration were contained within a single Snakemake v5.18.1 (Köster & Rahmann, 2018) script.

We used VCFtools v0.1.16 (Danecek et al., 2011) for additional rounds of filtering. First, we removed low quality sites with quality score <25, read depth <12, and non-biallelic sites. Second, we removed sites and individuals with a genotyping rate less than 50%. Third, we removed all sites that were on unresolved haplotigs by retaining only those SNVs that were on one of seven autosomal scaffolds (GenBank Nucleotide accessions: HE601624.2-30.2), the sex-linked ZW scaffold (HE601631.2), or the mitochondria (HE601612.2). Finally, for analyses requiring unlinked SNVs, we filtered linked sites within 250Kb windows using Plink v1.90b4 (Purcell et al., 2007) with the following parameters “— indep-pairwise 250kb 1 0.20”.

### Summary Statistics

We quantified read depth per probed-exome region with MosDepth v0.2.5 (Pedersen & Quinlan, 2018) and calculated genome-wide summary statistics for each population, including F_3_, F_ST_, Tajima’s *D*, π, and the Watterson estimator (Θ) with scikit-allele v1.2.1 (Miles, Ralph, Rae, & Pisupati, 2019). We examined genome regions that were targeted by the Le Clec’h *et al*. (2018) probe set for these calculations, non-target regions (i.e. non-exomic) were ignored since most samples lacked information from these regions. F_ST_ between populations was calculated form the average Weir-Cockerham F_ST_ (Weir & Cockerham, 1984) in windows of 100 SNVs. Effective population size (*Ne*) was estimated from Θ and the mutation rate (μ = 8.1 e-9 per base per generation; Crellen et al., 2016) with:

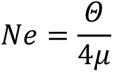

We examined linkage disequilibrium (LD) within each population by calculating *r*^2^ (--r2) values with PLINK v1.90b6.18 (Purcell et al., 2007). We excluded invariant sites from the analyses. Intra-autosomal, pairwise comparisons between SNVs within 1Mb of one another were allowed by setting the following parameters: “--ldwindow 1000000”, “--ld-window-kb 1000”, and “--ld-window-r2 to 0.0”. *r*^2^ values were then binned into 500 bp windows and averaged for each population using the R v3.6.1 stats.bin function in the fields v11.6 (Nychka, Furrer, Paige, & Sain, 2017) library. We used local regression to smooth the binned *r*^2^ values with the loessMod function in the base R v3.6.1 package and a span size of 0.5.

We used a Pearson Mantel test to examine correlation between genetic and physical distance. Since we did not have exact collection coordinates from whole genome samples, or they were lab derived, we excluded them from the analyses and instead focused only on the *S. mansoni* exome samples. We calculated pairwise p-distances with VCF2Dis (https://github.com/BGI-shenzhen/VCF2Dis; commit: b7684d3, accessed 13 Feb 2021) and physical distances between samples the Python haversine 2.3.0 module. Finally, we used the mantel() function in the scikit-bio 0.2.1 Python library to conduct a Pearson Mantel test that included 1,000 permutations.

### Population structure and admixture

We examined population substructure using PCA and Admixture with unlinked autosomal SNVs (described above). Two PCAs were calculated in PLINK v1.90b6.18, with and without the *S. rodhaini* samples. Population ancestry was estimated with ADMIXTURE v1.3.0 (Alexander, Novembre, & Lange, 2009). We examined between *k*=1 to *k*=20 populations and used the Cross-validation scores and the Evanno *et al*. (2005) method were used to determine a range of viable *k*’s. Q estimates were used as proxy for ancestry fractions.

We examined D (Patterson et al., 2012), D_3_ (Hahn & Hibbins, 2019) and F_3_ (Patterson et al., 2012) to identify gene flow between *S. rodhaini* and *S. mansoni* populations with emphasis on the Tanzanian *S. mansoni* population since it is from East Africa as are the *S. rodhaini* samples. For D_3_, we calculated the mean-pairwise (Euclidean) distances between populations using scikit-allele’s allel.pairwise_distance() function. To determine significance, we used 1,000 block bootstrap replicates of 1,000 SNV blocks. We calculated the average F3 across the genome in blocks of 100 variants. Here we ran multiple test that included some combination of an African *S. mansoni* population (Niger, Senegal, and Tanzania) as the test group and Brazilian *S. mansoni* and *S. rodhaini* as the potential source populations. D, or the ABBA-BABBA statistic, averaged over blocks of 1,000 variants assuming a phylogeny of (((*a*, Tanzania), *S. rodhaini*), *S. margrebowiei*), where the *a* population was either Brazil, Niger, or Senegal. D, D_3_ and F_3_ values were calculated using scikit-allel.

### Phylogenetics

We used three different phylogenetic methods to visualize relationships among sampled schistosomes: a mitochondrial haplotype network, a coalescent-based species tree, and a phylogenetic network.

(i) <u>Mitochondrial haplotype network</u>. We extracted mitochondrial SNVs from all *S. mansoni* individuals with VCFtools and converted the subsequent VCF file to Nexus format with vcf2phylip v2.0 (Ortiz, 2019). A median joining network (ε = 0) was created in from the mitochondrial haplotypes with PopArt v1.7 (Leigh & Bryant, 2015).
(ii) <u>Coalescent-based species tree</u>. We generated a coalescent-based species tree with SVDQuartets (Chifman & Kubatko, 2015) packaged in PAUP* v4.0.a.build166 (Swofford, 2003). We examined parsimony-informative, autosomal SNVs by removing private alleles (singleton and doubletons). All samples were assigned to a population based on their country of origin (ex. Niger, Puerto Rico, Brazil, Cameroon, etc.) except for the lab-derived, Caribbean samples. Each of these samples was considered to represent an individual population given their histories of extensive lab passage (ex. Guadeloupe1, Guadeloupe2, Puerto Rico). We randomly evaluated 100,000 random quartets and bootstrapped the quartet tree with 1,000 standard replicates. The tree was rooted on the single *S. margrebowiei* individual.
(iii) <u>Phylogenetic network</u>. We used a phylogenetic network to visualize and quantify migration among schistosome populations. We only included *S. mansoni* populations with more than 4 individuals, which excluded all whole genome samples from this analysis including those from the Caribbean and the *S. rodhaini* samples. We used autosomal SNVs after filtering linked sites in 250Kb blocks with PLINK v1.90b6.18 and then used TreeMix v1.12 (Pickrell & Pritchard, 2012) to generate the phylogenetic network. This analysis used a co-variance matrix generated from blocks of 500 SNVs without sample-size correction (“--noss”) and the number of migration events was limited to 3.

### Selection

We scanned the genome to identify regions under selection using haplotype (H-scan v1.3; Schlamp et al., 2016), allele-frequency (SweepFinder2 v2.1; DeGiorgio, Huber, Hubisz, Hellmann, & Nielsen, 2016), and PCA-based (pcadapt v4.3.3; Luu, Bazin, & Blum, 2017) methods. In addition, to avoid false positives, we used msprime v0.7.4 (Kelleher, Etheridge, & McVean, 2016) to conduct simulations to estimate the range of values expected under neutrality from H-Scan and SweepFider2.

(i) <u>Neutral simulations</u>. We used msprime v0.7.4 (Kelleher et al., 2016) to simulate a set of neutrally evolving SNVs along a single chromosome for each population and then used the simulated data with H-Scan and SweepFinder2 to define the range of values expected in the absence of selection. For these simulations we used a mutation rate (μ = 8.1 e-9 per base per generation; Crellen et al., 2016) and recombination rate (3.4e-8 per base per generation; Criscione, Valentim, Hirai, LoVerde, & Anderson, 2009) from previous work on *S. mansoni*. Population-specific estimates of *Ne* are described above. The chromosome length was set to 88.9 Mb which is equal to chromosome 1 (HE601624.2) in the *S. mansoni* assembly. The number of chromosomes sampled per msprime run was equal to the number of samples we collected in each population. We used these parameters to perform 342 simulations for each population, roughly equivalent to 100 genomes worth of simulated data. Next, we down sampled the SNVs along the entire simulated chromosome so that they were comparable with our targeted sequencing approach (i.e. only SNVs from “exomic” regions). Since the simulated chromosomes were the same size as chromosome 1 we transposed the chr1 annotation onto the simulated chromosome and extracting only those SNVs occurring in regions accessible by our biotinylated probes with VCFtools. The simulated data was run through H-Scan and SweepFinder2 in parallel with the actual SNV data to establish a range of neutral values for each test (described above).
(ii) <u>H-scan</u>. This method measures the length of homozygous haplotypes to identify regions under selection. Strong selection on adaptive alleles drives SNVs under selection and any linked alleles to high frequency and reduces homozygosity in the surrounding region. For each population, we converted autosomal SNVs from VCF to H-Scan format using vcf2hscan.py script from vcf2phylip v2.0 (Ortiz, 2019) and ran H-Scan on each chromosome with a maximum gap length (-g) of 10Kb. *H* values were smoothed (*H_smoothed_*) by median filtering values in 201 SNV windows (step size = 1) using the medfilt function in the SciPy v1.5.2 (Virtanen et al., 2020) for visualization purposes.
(iii) <u>SweepFinder2</u>. This method uses deviations in allele frequency from a neutral expectation to estimate the selection while accounting for the possibility of background selection via a likelihood ratio (LR) test (DeGiorgio et al., 2016). Empirical site-frequency spectra were calculated for each population and within each population LR was estimated along each autosome individually. We examined grid points (‘g’), or window sizes, of 1, 5, 10, and 20 Kb with minimal impact on the results. Downstream analyses are reported on the runs with ‘g’ = 1kb.
(iv) <u>*pcadapt*</u> – We used the R v4.0.5 package pcadapt v4.3.3 (Luu et al., 2017) to identify highly differentiated loci among populations via variants associated with population structure as identified by PCA. We only included all samples from Brazil, Niger, and Senegal since our primary goal was to identify variants involved in adaptation the Americas. Rare variants (MAF<5%) were excluded with VCFtools v0.1.16. We identified the appropriate number of principal components from the data by running an initial *pcadapt* run with 20 populations (K=20) and LD filtered variants (LD.clumping = list(size = 100, thr = 0.2)). The major break in the subsequent scree plot was used as the optimal K choice. We used a second pcadapt run with the optimal K and the same LD filtering parameters as the initial run to assign *p* values to each site. Finally, we adjusted *p* values for multiple tests with Bonferroni correction and an α = 0.05 to identify SNV outliers associated with population differentiation.
(v) <u>Identifying regions of selection-</u> We identified regions potentially under positive selection using a three-step process. First, we identified SNVs whose H-Scan and Sweepfinder2 values were in the 99^th^ percentile of and greater than the neutral thresholds established with msprime. These were SNVs with the strongest signal of selection. Then, we expanded from the SNV to a broader region by merging all variants within 333,333 bp whose H-Scan or Sweepfinder2 values were greater than the neutral thresholds. Finally, we looked for pcadapt outliers in each region. These regions are referred to as “putative selected regions” or “putative regions of selection”. Gene names and putative functions were taken from UniProtKB (release 2020_06) or HHsearch (Steinegger et al., 2019) annotations from Le Clec’h et al. (2021). The entire process is summarized in Figure 1.

**Figure 1.**
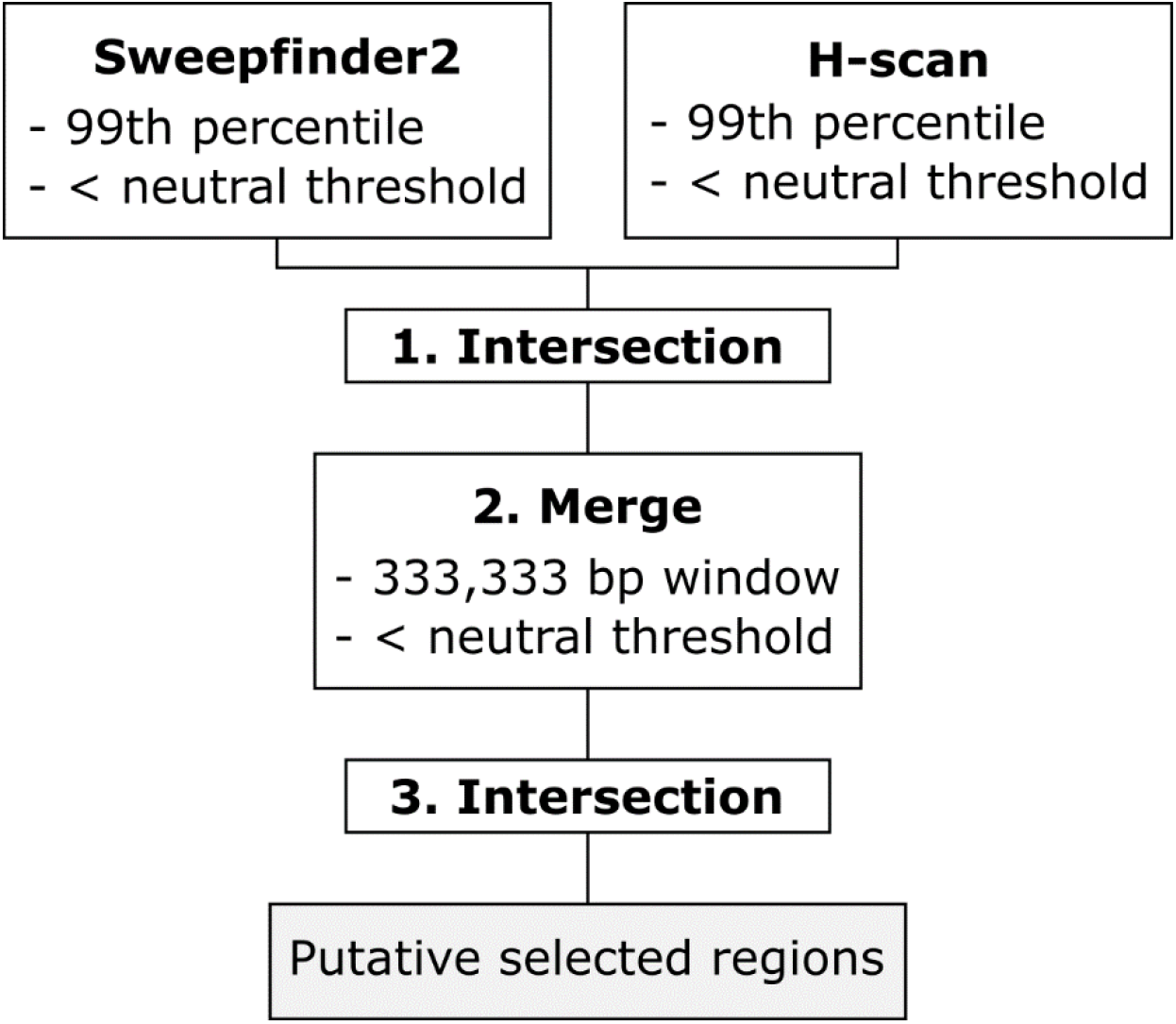
Identifying regions of selection. Flow chart shows how three separate analyses were combined to identify regions most likely experiencing positive selection. These regions are referred to as “putative selected regions”. We established neutral thresholds by simulating neutrally evolving SNVs for each population. The simulated data was then run through H-Scan and Sweepfinder2 to determine the maximum expected values from neutral data for each analysis.

## Results

### Summary of Sequence Data

After genotyping and filtering, we removed 25 of the 178 samples with low numbers of reads, poor coverage, or low genotyping rates. The final dataset included 135 *S. mansoni* (exome), 8 *S. mansoni* (genome), 8 *S. rodhaini* (exome), 1 *S. rodhaini* (genome), and 1 *S. margrebowiei* (genome). We genotyped 1,823,890 sites, which was reduced to 475,081 autosomal and 815 mitochondrial variants after quality filtering. The final dataset comprises 153 samples with mean read depths of 520.7x (range: 251.4-998.2x) and 66.0x (range: 14.8-726.2) at mitochondrial and autosomal loci. Location and sequence coverage statistics for all samples in the final dataset are listed in Supplemental Table 1.

### Summary statistics

Autosomal and mitochondrial summary statistics for π, *H*, Tajima’s *D*, Θ, and *Ne* are shown in Table 1. π, Θ, and *Ne* are similar between the West African and Brazilian *S. mansoni* populations but are 2-3 times lower than observed in Tanzanian *S. mansoni*. Tajima’s *D* values range between slightly positive to negative (Tajima’s *D* = -1.417 – 0.034) in *S. mansoni* (Table 1; Supplemental Table 2). All the African populations show negative Tajima’s *D*, consistent with natural selection or population expansion. However, the Brazilian Tajima’s *D* is positive, which is inconsistent with a bottleneck during colonization of South America. Mitochondrial diversity was quantified with π and haplotype diversity (*H*). *H* in all populations is very high (<0.978) indicating that all mitochondrial haplotypes are unique. Mitochondrial π follows the same pattern as autosomal π with the exception that the mitochondrial π in Brazil is lower than may be expected when compared to measures in Niger and Senegal. F_ST_ values between *S. mansoni* populations are shown in Table 2 and were highest in pairwise comparisons that included the Tanzanian population. Mantel tests showed significant signs of isolation-by-distance within Africa (*r* = 0.64, *p* = 0.001) and in African and Brazilian (*r* = 0.77, *p* = 0.001) *S. mansoni* samples.

**Table 1.** Whole genome summary statisitics for *Schistosoma mansoni* populations and *S. rodhaini*

|  | **n** | **$\pi$** | **$\pi$ (mito)** | **$H$ (mito)** | **Tajima's $D$** | **$\Theta$** | **$N_e$** | **$\pi$ (PRS)** | **Tajima's $D$ (PRS)** |
| --- | --- | --- | --- | --- | --- | --- | --- | --- | --- |
| *S. rodhaini* | 9<sup>a</sup> | 5.61E-04 | Na<sup>b</sup> | Na<sup>b</sup> | 0.479 | 4.29E-04 | 13,226 | Na<sup>c</sup> | Na<sup>c</sup> |
| Sm (Brazil) | 45 | 6.93E-04 | 2.10E-03 | 0.984 | 0.034 | 5.84E-04 | 18,032 | 2.22E-04 | -1.218 |
| Sm (Niger) | 10 | 6.00E-04 | 5.89E-03 | 0.978 | -0.579 | 6.07E-04 | 18,737 | 2.46E-04 | -1.018 |
| Sm (Senegal) | 25 | 4.97E-04 | 4.72E-03 | 0.997 | -1.417 | -1.375 | 21,992 | 1.75E-04 | -1.816 |
| Sm (Tanzania) | 55 | 1.45E-03 | 7.25E-03 | 1.0 | -0.729 | -0.739 | 51,508 | Na<sup>c</sup> | Na<sup>c</sup> |

"Sm" - *Schistosoma mansoni*, "n" - number of samples, " $\pi$ " - nucleotide diversity, " $H$ " - haplotype diversity, " $\Theta$ " - Watterson estimator, " $N_e$ " - effective population size, "PRS" - putative region of selection
a) 8 of 9 *S. rodhaini* samples came from a single lab population: population statistics are likely biased
b) No *S. rodhaini* reads mapped to the *S. mansoni* mitochondria.
c) Not calculated.

**Table 2.**
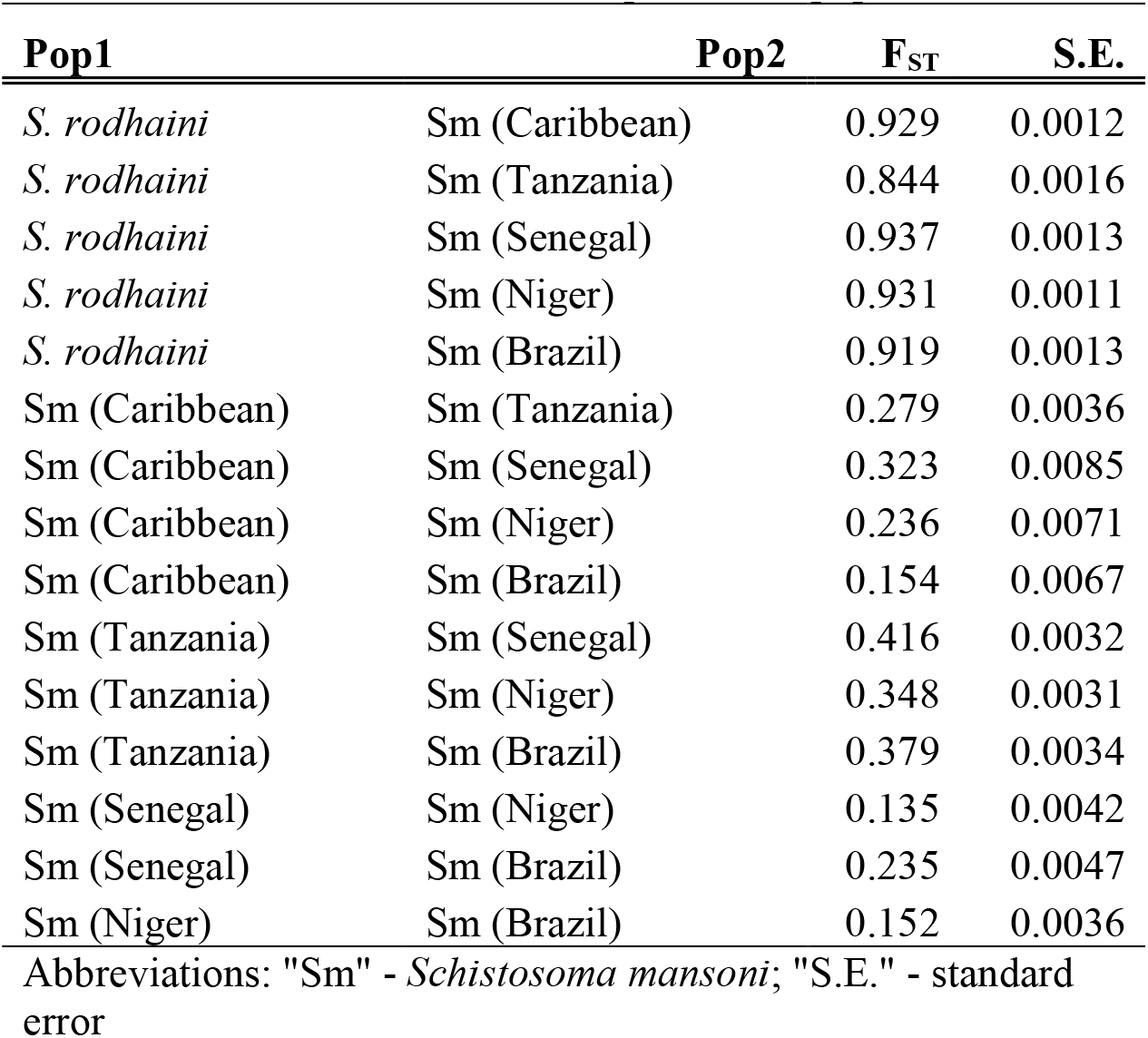
F_ST_ between Schistosoma species and populations

| **Pop1** | **Pop2** | **$F_{ST}$** | **S.E.** |
| --- | --- | --- | --- |
| *S. rodhaini* | Sm (Caribbean) | 0.929 | 0.0012 |
| *S. rodhaini* | Sm (Tanzania) | 0.844 | 0.0016 |
| *S. rodhaini* | Sm (Senegal) | 0.937 | 0.0013 |
| *S. rodhaini* | Sm (Niger) | 0.931 | 0.0011 |
| *S. rodhaini* | Sm (Brazil) | 0.919 | 0.0013 |
| Sm (Caribbean) | Sm (Tanzania) | 0.279 | 0.0036 |
| Sm (Caribbean) | Sm (Senegal) | 0.323 | 0.0085 |
| Sm (Caribbean) | Sm (Niger) | 0.236 | 0.0071 |
| Sm (Caribbean) | Sm (Brazil) | 0.154 | 0.0067 |
| Sm (Tanzania) | Sm (Senegal) | 0.416 | 0.0032 |
| Sm (Tanzania) | Sm (Niger) | 0.348 | 0.0031 |
| Sm (Tanzania) | Sm (Brazil) | 0.379 | 0.0034 |
| Sm (Senegal) | Sm (Niger) | 0.135 | 0.0042 |
| Sm (Senegal) | Sm (Brazil) | 0.235 | 0.0047 |
| Sm (Niger) | Sm (Brazil) | 0.152 | 0.0036 |

Abbreviations: "Sm" - *Schistosoma mansoni*; "S.E." - standard error

Diversity and *Ne* values were lower in *S. rodhaini* than all *S. mansoni* populations but should be interpreted with caution. Eight of the nine *S. rodhaini* samples came the same laboratory-maintained population. F_ST_ between *S*. *rodhaini* and individual *S. mansoni* populations ranged from 0.844 (*S. rodhaini* vs. Sm Tanzania) - 0.937 (*S. rodhaini* vs. Sm Senegal).

Figure 3 shows linkage disequilibrium (LD) decay (binned and smoothed) from pairwise *r*^2^ values. LD decays to *r*^2^ ≤ 0.2 within 500Kb for all populations. LD was weakest in the Tanzania population with *r*^2^ decaying to 0.5 in 28 bp. LD decay in the other three populations (Senegal, Niger, and Brazil) was relatively consistent with LD decaying by half (*r*^2^ = 0.5) between 15,150 bp (Niger) and 26,196 bp (Brazil). Both Senegal and Niger show high levels of LD even between SNVs on different chromosomes: this results from lower sample size (Senegal, n=25; Niger, n=10) in these two populations.

**Figure 3.**
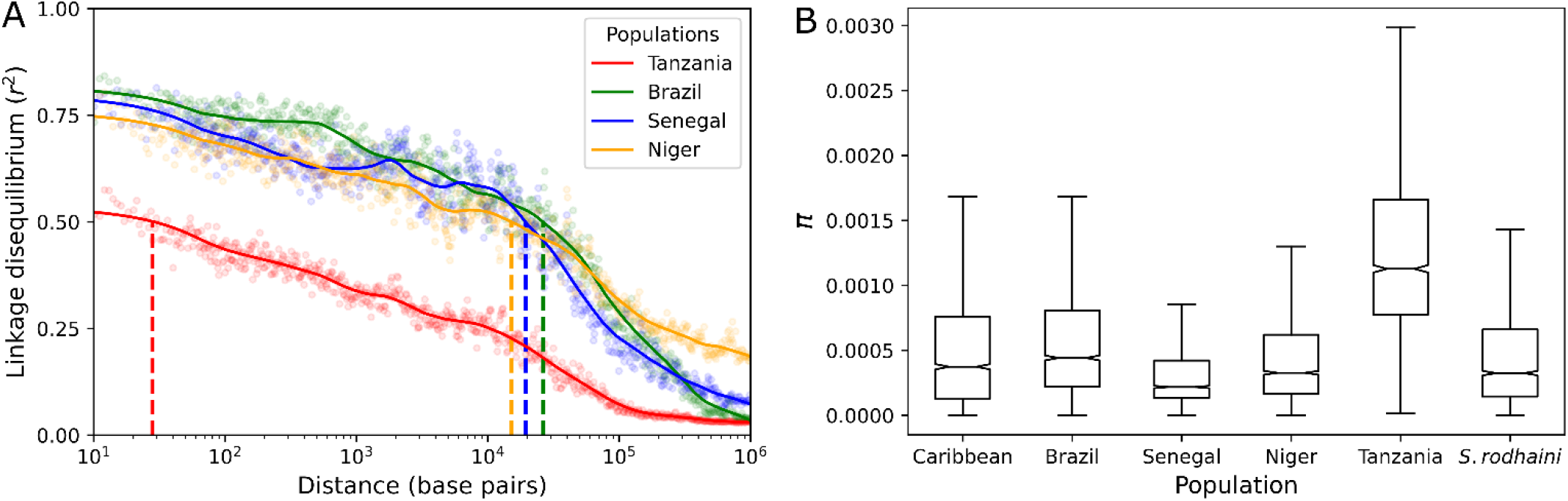
Linkage Disequilibrium Decay and diversity within populations. (A) Linkage disequilibrium between SNVs was quantified with *r*^2^ values for each population. Mean *r*^2^ values were taken in 500 bp windows and loess smoothed. Vertical dotted lines indicate the distance where *r*^2^ = 0.5 for each population. LD decayed to *r*^2^=0.5 in 28 bp (Tanzania), 15,150 bp (Niger), 19,318 bp (Senegal), and 26,196 bp (Brazil). (B) Nucleotide diversity (π) varied between *S. mansoni* populations with the highest levels of diversity occurring in E. Africa (Tanzania). Π was measured in 100 Kb windows across the autosomal chromosomes. Outliers are not shown.

### Admixture with S. rodhaini

We asked whether hybridization with *S. rodhaini*, a closely-related schistosome infecting rodents, might contribute to the high genetic diversity observed in East Africa vs West Africa/south American *S. mansoni*. To investigate this, we used three statistics (D, D_3_, and F_3_) to test for admixture between *S. mansoni* and *S. rodhaini*, with particular emphasis on the Tanzanian populations of *S. mansoni*. Each of these statistics, attempts to identify the presence of admixture in different ways. D and D_3_ values ≠ 0 indicate admixture and positive and negative values determining the direction of introgression. F_3_ values < 0 indicate admixture between the two source populations. None of the three statistics, or any of the population combinations, returned values containing significant signals for admixture (Table 3). These results suggest that hybridization/introgression between *S. mansoni* and *S. rodhaini* may make no detectable contribution to elevated diversity in East Africa.

**Table 3.**
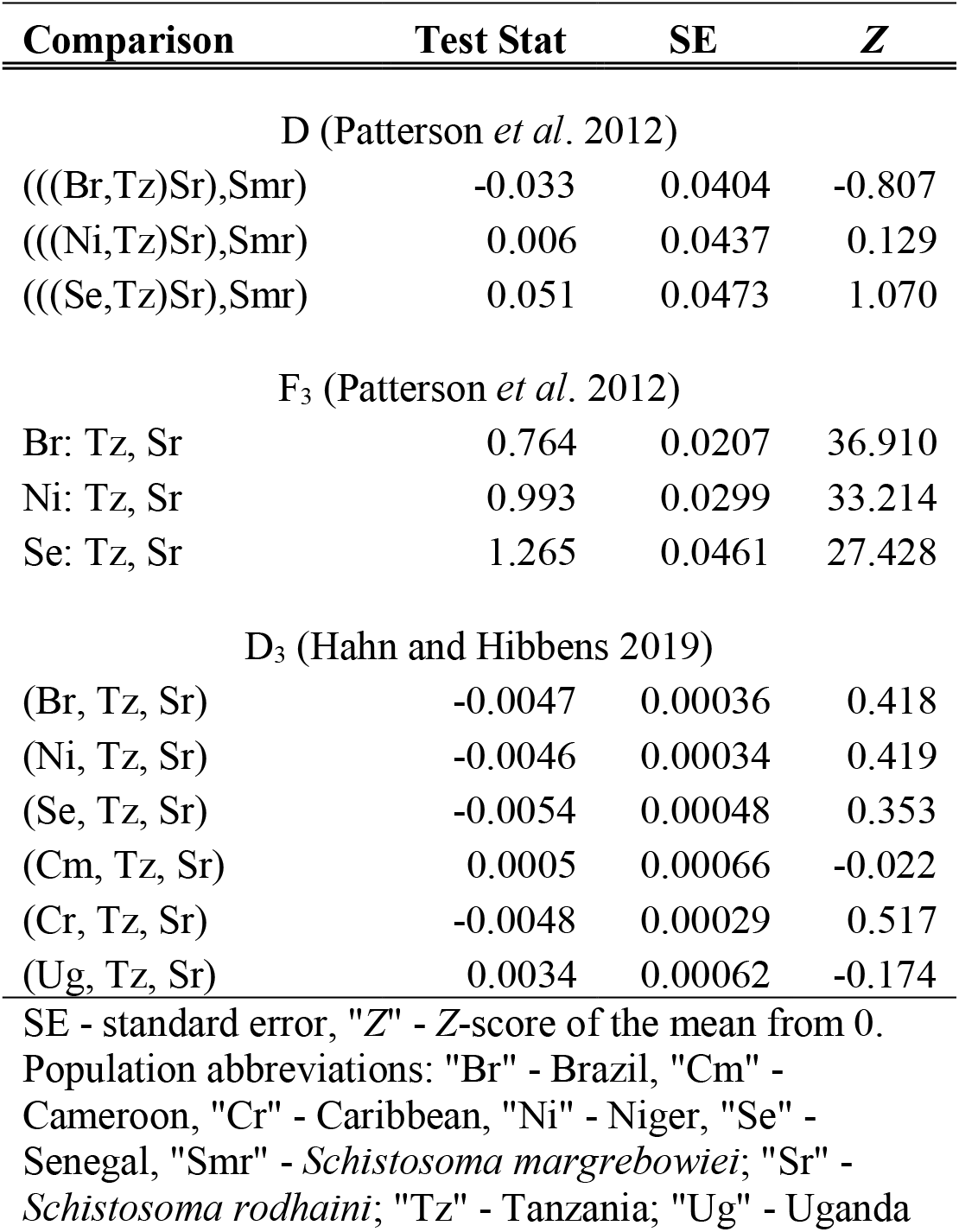
Admixture statistics

### Population structure

We examined population structure using PCA and ADMIXTURE with 38,197 unlinked autosomal SNVs. Two PCAs were generated, with and without the *S*. *rodhaini* outgroup (Figure 4). The two species were differentiated along PC1 (34.7% variance) when *S*. *rodhaini* was included (Figure 4A). *S. mansoni* samples cluster into geographically defined groups when *S. rodhaini* is excluded (Figure 4B). East African samples were distinct from all other *S. mansoni* samples, except for one whole genome sample collected in Kenya (see below). Samples from the Americas, including those from the Caribbean and Brazil, showed a closer relationship with Cameroon and Nigerien samples than those from Senegal.

**Figure 4.**
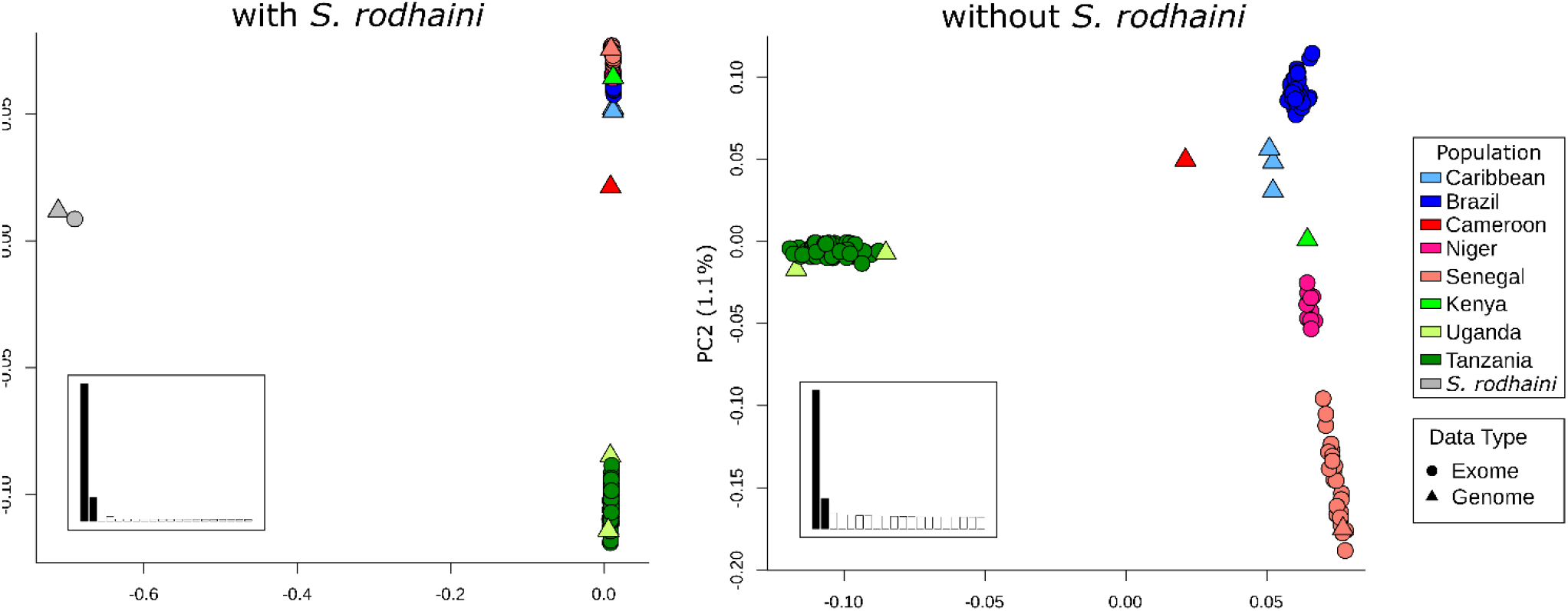
A PCA of unlinked autosomal SNVs. The PCA plot that included *S*. *mansoni* and *S. rodhaini* (left) clearly shows a large distinction between the two species with some variation within *S*. *mansoni* along PC2. A PCA with only *S. mansoni* (right) differentiates East African *S. mansoni* along PC1. The remaining *S. mansoni* samples fall along a continuum on PC2 that goes from samples in West Africa and transitions to the Americas.

We used ADMIXTURE to assign individuals to one of *k* populations, where *k* is between 1 and 20 (Figure 5). Cross-validation scores (Evanno et al., 2005) were minimized when *k* was 4 or 5. Both *k* = 4 or 5 split *S. mansoni* samples into geographically defined populations with two major differences. First, *k*=4 showed that the allelic component primarily associated with Brazil was found at moderate levels in Cameroon and Nigerien individuals. Second *k*=5 split the West African samples into a Senegalese and a Cameroonian + Nigerien populations.

**Figure 5.**
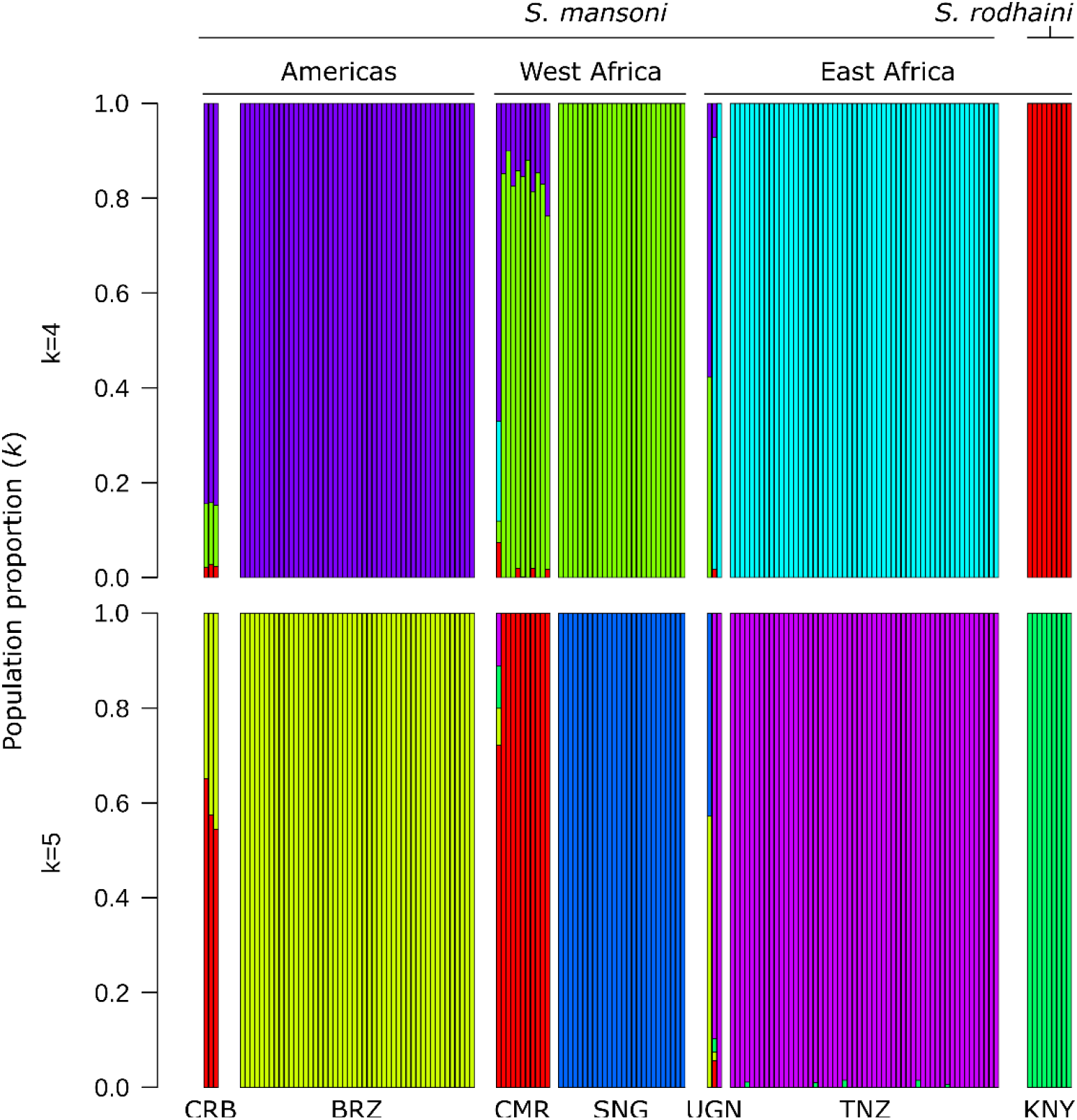
**Population structure in *S. mansoni*** ADMIXTURE analyses with k=4 and k=5 populations identified clear distinctions between each of the major sampling localities (the Americas, West Africa, East Africa, and *S*. *rodhaini*). The population components in each of the whole genome samples from Uganda (UGN), Kenya (KNY), Cameroon (CMR), and the Caribbean (CRB) were more heterogeneous than samples with exome data. Cameroonian and Nigerien samples contain moderate proportions of the Brazilian population component. Abbreviations: “BRZ” – Brazil, “SNG” – Senegal, “NGR” – Niger, “TNZ” – Tanzania, “BRN” – Burundi.

As observed in the PCA, the Kenyan , whole-genome sample contained a large portion of alleles associated with samples from the Americas (∼40-60%) in the ADMIXTURE analysis. Crellen *et al*. (2016) recovered similar results and hypothesized that the Kenyan sample may be reflecting human-trafficking routes between Portuguese and Arab slave traders out of the port of Mombasa. Given that only a single sample is available from this region, and contamination with South American strains during laboratory passage is a possible alternative explanation, we chose to remove the Kenyan sample from downstream analyses.

### Phylogenetics

We used three different phylogenetic methods to investigate the evolutionary relationships between sequences (Figure 6). First, we generated median-joining network from 815 mitochondrial SNVs of which 477 where phylogenetically informative (Figure 6A). The haplotype network identified three major haplotypes roughly corresponding with the geographic partitioning of the samples. The haplogroups include an East African clade (Tanzania, Uganda), a Senegal group, and intermediate haplogroup with samples from the Brazilian and Nigerien populations. Caribbean samples were not assigned to a single haplogroup. The single sample from Puerto Rico was associated with the major Brazilian and Nigerien haplotype, and the two Guadalupe samples tended to be more strongly associated with Senegalese haplotypes. Additionally, the sample from Cameroon was only single step removed from the most common Brazilian haplotype.

**Figure 6.**
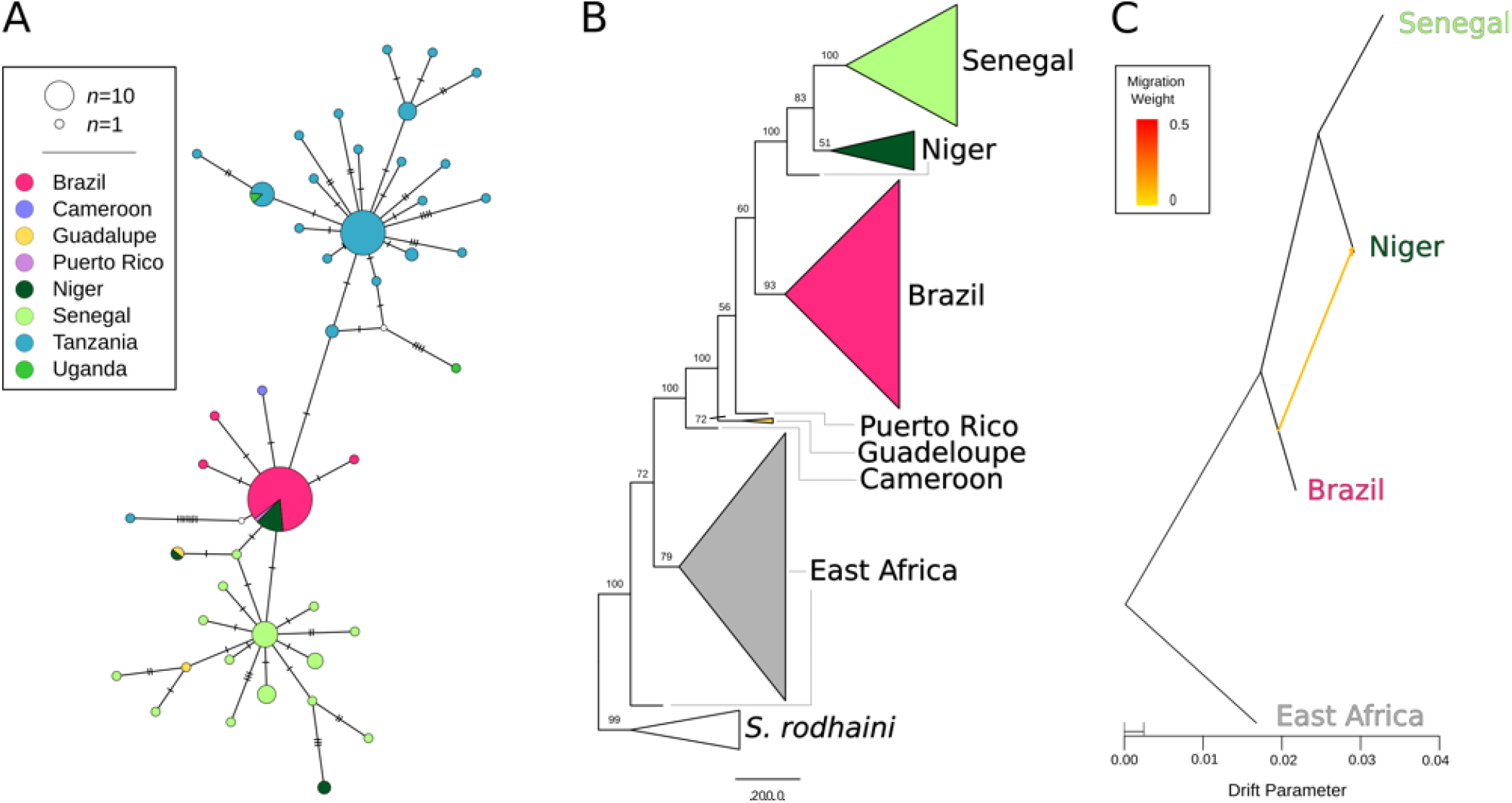
Phylogenetic relationships between *S. mansoni* populations. Multiple phylogenetic analyses and marker types were used to discern relationships between *S. mansoni* populations. (A) A median-joining haplotype network was constructed from 815 variations variants across the mitochondria of all *S. mansoni* samples. (B) A coalescent-based species tree from 100,819 parsimony informative with bootstrap values shown on each clade. Monophyletic populations are shown as a collapsed clade except in the case of E. Africa which contains samples from Tanzania and Uganda. (C) A maximum likelihood phylogenetic network of autosomal variants identified a single, weak migration edge oriented from Brazil to Niger. All three analyses identify a relationship between Senegal, Niger, and Brazil that excludes East African samples. The mitochondrial (A) and autosomal (C) networks both allow for direct relationship or allele sharing between Brazil and Niger. The species tree (B) indicates a strong relationship between Senegal and Niger that excludes Brazil (bootstrap percentage = 100).

A coalescent-based species tree from 100,819 parsimony informative SNVs was generated with SVD-quartets (Figure 6B). Quartet sampling was limited to 100k quartets which sampled 0.43% of all distinct quartets present in the alignment. The final species tree was consistent with 84.7% of all the quartets sampled. Unlike the mitochondrial tree, samples fall into well supported clades corresponding with geography with two exceptions. Samples from East Africa and Niger formed independent paraphyletic clades. In both cases paraphyly was induced by a single individual. Bootstrap support was generally higher in the quartet species tree than in the mitochondrial tree. West African samples formed a well-supported monophyletic clade with the Brazilian and Caribbean samples, indicating a shared origin for these parasites. Brazilian and Caribbean samples appear to have a polyphyletic origin within a larger clade containing West African parasites.

Finally, migration between populations was quantified with TreeMix (Figure 6C). We only examined populations with more than 5 individuals which excluded Cameroonian and Caribbean samples from the analysis. The topologies linking the remaining populations (East Africa, Niger, Senegal, and Brazil) in the TreeMix and species tree were identical. The species tree was slightly improved with the addition of a single migration edge from Brazil to Niger. This migration edge significantly improved the likelihood score of the topologies from 73.6026 to 73.7165 with an edge weight of 0.1 (*p* = 0.0081).

*Selection* – We used msprime to generate a set of neutrally evolving SNVs based on parameters specific to each of the sampled populations. These neutrally evolving SNVs were distributed across an 88.9 Mb chromosome that was equal in size to *S. mansoni* chromosome 1 (HE601624.2). We then transposed the HE601624.2 exome annotation onto the simulated chromosome to extract “exome” data. This process was repeated 342 times per population to produce a set of neutrally evolving loci to use as controls when examining selection on actual samples. We used these neutral simulations to generate maximum (100%) threshold values for H-Scan and Sweepfinder2 test statistics expected under neutrality (see below). The mean 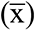 simulated SNV count across all replicates was: 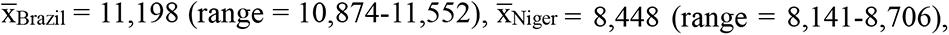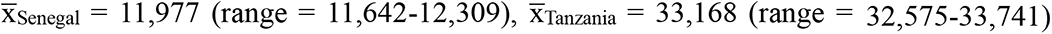. The number of observed SNVs was between 72.2-89.7% of the mean number of simulated SNVs (Brazil = 8,947; Niger = 6,107; Senegal = 9,381; Tanzania = 29,765). The reduced number of SNVs in the neutral data is likely due to the absence of selection that is presumed to be acting on the exome data sets.

The H-Scan and SweepFinder2 results are shown in Figure 7. *pcadapt* results are show in Supplemental Figure 1. Each program uses different methodology to detect sites under selection. H-Scan calculates *H* with homozygous tract length and the number of haplotypes to identify genome regions that have undergone selective sweeps. *H* values were highly variable for each population, even within windows smaller than 100Kb. In Tanzania only 2 of the 475,081 SNVs had *H* values higher than were generated from neutral simulations. SweepFinder2 calculates deviations from a neutral site frequency spectrum correcting for the possibility of background selection. A likelihood ratio (LR) describes the probability of positive selection vs. neutral evolution and background selection within a designated window. Sweepfinder2 was able to clearly define multiple peaks for each population when compared to H-Scan. Each of the four populations had regions greater than neutral expectations. *pcadapt* identifies SNVs significantly associated with population differentiation. In this analysis we found 442 SNV outliers, after multiple test correction, that were distributed across all seven autosomes at 127 loci. These regions were distributed across 280.2 Mb. F_ST_ of the *pcadapt* outliers (F_ST_ = 0.544) was significantly higher than in the remaining population outliers (F_ST_ = 0.195).

**Figure 7.**
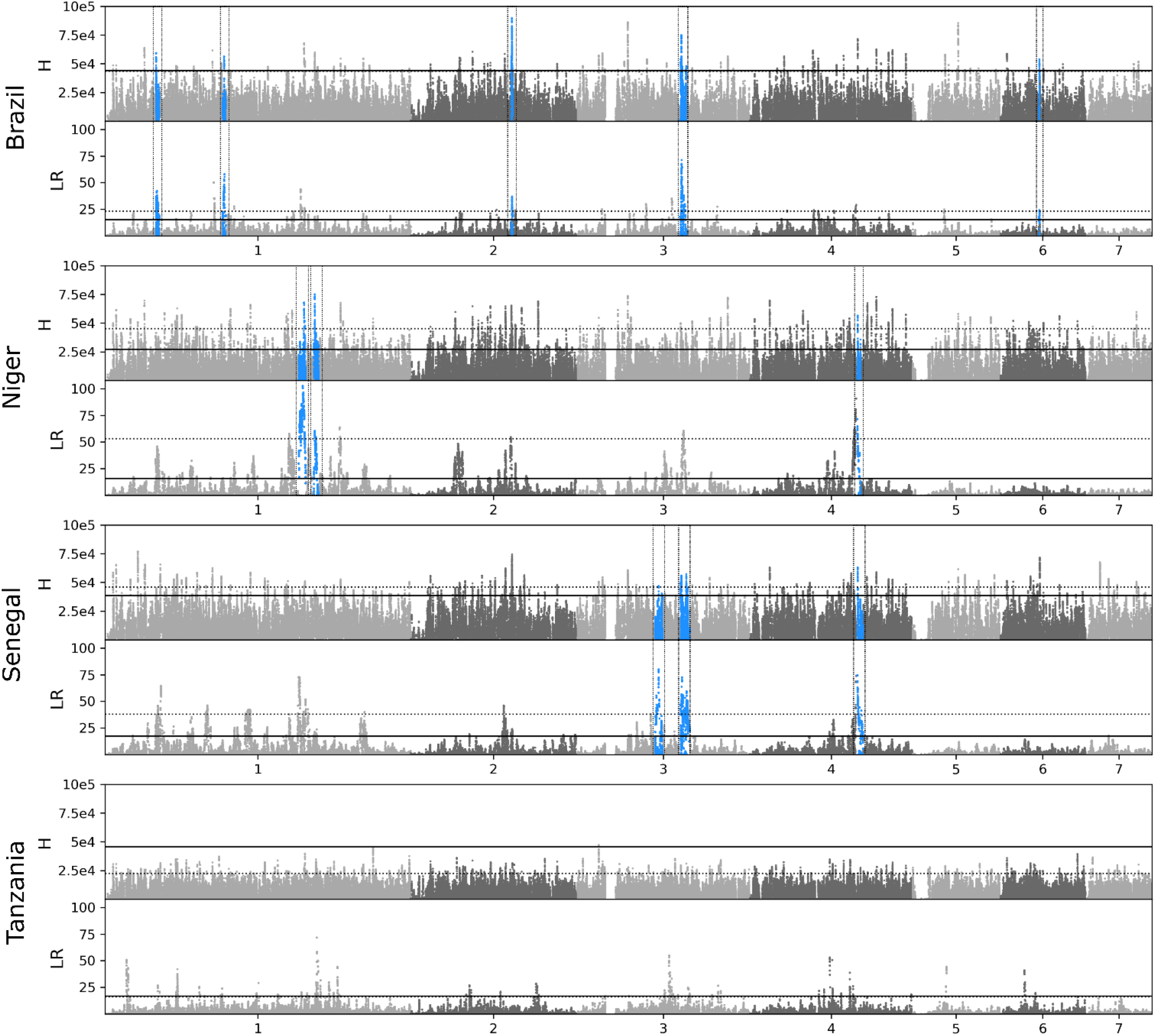
Positive Selection across the *S. mansoni* genome. Selection across the *S. mansoni* genome was calculated with haplotype (*H*) and allele frequency (likelihood ration; LR) - based methods. Dotted lines represent positions with an *H* or LR value in the 99^th^ percentile. The solid black line represents the maximum *H* or LR calculated form simulated data under neutral conditions. Regions of interest (blue, boxed) were identified by finding sites where the *H* and LR values were in the >99^th^ percentile and were both greater than the max *H* or LR from simulated data. Once these sites were identified we combined variants within 333,333 bp windows that showed signs of selection; an *H* or LR greater than the simulated threshold.

We defined “putative regions of selection”, as those that have most likely experienced positive selection. These regions contain variants (i) with both *H* and LR values in the 99^th^ percentile (ii) are greater than the neutral thresholds and (iii) have a signal of population-specific directional. All SNVs meeting one or more of these criteria are listed in Supplemental Table 4.

Our results recovered 5, 3, and 3 putative selected regions in Brazil, Niger, and Senegal respectively (Figure 7; Supplemental Table 5). Information regarding the number of regions, SNVs, and genes identified are presented in Table 4 and Table 5. π (Supplemental Figure 2) and Tajima’s *D* (Supplemental Figure 3) were depressed in these regions compared to genome-wide values (Supplemental Table 3) which is consistent with loci experiencing selection. On average the size of each region was relatively small (1,395,643 bp) and in two instances, these sites were shared between populations. The Brazilian and Senegalese populations shared a site on chromosome 3 at HE601626.2:30,092,830:31,936,551, while the Nigerien and Senegalese populations shared a peak on chromosome 4 at HE601627.2:31,216,154:32,138,352. We did not recover any regions or sites of interest in the Tanzanian population in large part because only two of 475,081 sites had higher *H* values the largest *H* from neutral simulations (*H* = 45532.9). These variants are on chromosome 3 at HE601626.2:6,500,422 and HE601626.2:6,503,809 and are adjacent to each other in our filtered SNV dataset.

**Table 4.**
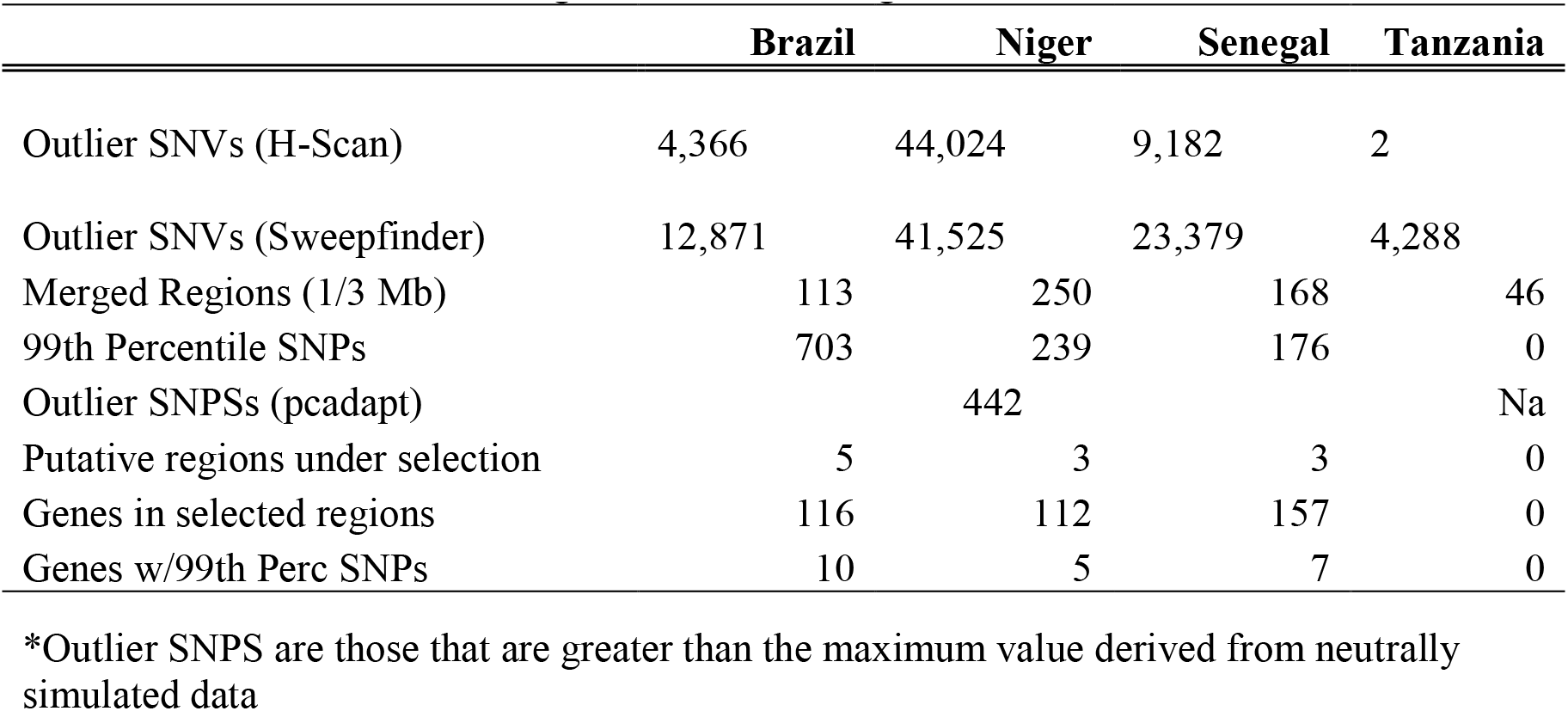
Number of SNPs and regions indentified in genome-wide scans for selection

**Table 5.** Genes in *Schistosoma mansoni* populations with the strongest signals of directional selection

| Gene ID | UniProtKB Accession | UniProtKB Description | HHsearch Annotation |
| --- | --- | --- | --- |
| **Brazil** |  |  |  |
| Smp_060090 | G4VAJ9 | 40S ribosomal protein S12 | Ribosomal protein S12e Ribosome |
| Smp_073680 | G4V701 | Putative TATA-box binding protein | DNA-directed RNA polymerase II subunit |
| Smp_123510 | A0A5K4EKW3 | Vacuolar protein sorting-associated protein 16 homolog | Vps16_C |
| Smp_123520<sup>1</sup> | A0A3Q0KKK4 | Putative RNA M<sup>5</sup>U methyltransferase | rRNA (Uracil-5-)-methyltransferase Ruma |
| Smp_123570 | A0A3Q0KKM5 | BHLH domain-containing protein | Aryl hydrocarbon receptor nuclear translocator |
| Smp_123590 | A0A5K4EL22 | Uncharacterized protein | Swi5-dependent recombination DNA repair protein |
| Smp_148460 | A0A3Q0KPP3 | Putative neurofibromin | GAP related domain of neurofibromin |
| Smp_162000 | A0A5K4ETP0 | UBR-type domain-containing protein | E3_UbLigase_R4 |
| Smp_246630 | A0A5K4F427 | UBC core domain-containing protein | E2 Ubiquitin conjugating enzyme |
| Smp_341570 | A0A5K4FC62 | Uncharacterized protein | Uncharacterised protein family UPF0183 |
| **Niger** |  |  |  |
| Smp_008230 | G4V7H8 | Putative rab-18 | di-Ras2 |
| Smp_126620 | A0A3Q0KL42 | Uncharacterized protein | Ligand-binding domain of low-density lipoprotein receptor |
| Smp_165060 | A0A3Q0KRY3 | Uncharacterized protein | Uncharacterized protein |
| Smp_167890<sup>2</sup> | Q6BC90 | Peptide-methionine (R)-S-oxide reductase | C-terminal MsrB domain of methionine sulfoxide reductase PilB |
| Smp_313490<sup>2</sup> | A0A5K4F3H5 | Uncharacterized protein | Transforming protein RhoA, Rho-associated, coiled-coil |
| **Senegal** |  |  |  |
| Smp_070780 | G4VEM1 | UDP-glucose 4-epimerase | Uridine diphosphogalactose-4-epimerase |
| Smp_123440 | A0A3Q0KKL2 | Putative fad oxidoreductase | D-amino-acid oxidase |
| Smp_123520<sup>1</sup> | A0A3Q0KKK4 | Putative RNA M<sup>5</sup>U methyltransferase | rRNA (Uracil-5-)-methyltransferase Ruma |
| Smp_164560 | A0A3Q0KRR1 | Uncharacterized protein | Na |
| Smp_167890<sup>2</sup> | Q6BC90 | Peptide-methionine (R)-S-oxide reductase | C-terminal MsrB domain of methionine sulfoxide reductase PilB |
| Smp_213150 | A0A5K4EZI5 | Uncharacterized protein | Ribonucleases P/MRP protein subunit POP1 |
| Smp_313490<sup>2</sup> | A0A5K4F3H5 | Uncharacterized protein | Transforming protein RhoA, Rho-associated, coiled-coil |

1. Shared between the Brazil and Senegal
2. Shared between Niger and Senegal
UniProtKB descriptions are from release 2020\_06
HHsearch annotations are from Le Clec'h et al. (2021)

We identified 116-157 genes within “putative selected regions” in the Brazilian, Nigerien, and Senegalese populations (Supplemental Table 6). Within these populations 10, 5, and 7 genes contain SNVs meeting the 99^th^ percentile. Several genes identified in these regions were shared between populations. Brazil and Senegal shared 48 genes in target regions, and Senegal and Niger shared 22 genes in target regions. Three genes with 99^th^ percentile SNVs were shared between populations: Smp_313490 and Smp_167890 (Niger and Senegal), Smp_123520 (Brazil and Senegal; Table 5).

## Discussion

We examined the impact of human-mediated dispersal of *S. mansoni* during the Trans-Atlantic slave trade. Previous analyses with *S. mansoni* used mitochondrial data or had limited sampling from the Americas (Crellen et al., 2016; Morgan et al., 2005; Webster et al., 2013). To further these studies, we included 135 exome sequences available from natural populations in Brazil (n=45), Niger (n=10), Senegal (n=25), and Tanzania (n=55) which we combined with the existing genome sequences from Cameroon (n=1), the Caribbean (n=4), Senegal (n=1), and Uganda (n=2) (Berriman et al., 2009; Chevalier et al., 2016; Crellen et al., 2016). We used these data to examine *S. mansoni* expansion across Africa, explore examine parasite colonization of the Americas, quantify signatures of selection during colonization, and detect hybridization with *S. rodhaini*, a closely related parasite utilizing a rodent host.

### Elevated East African diversity and S. mansoni expansion across Africa

A striking result from this study is the dramatic reduction in genetic diversity between East and West Africa. Sequence summary statistics indicate that the East African population has 2-3-fold greater nucleotide diversity (π), larger *Ne* and greater mitochondrial diversity than the other populations (Table 1; Figure 6A). Phylogenetic analyses rooted with *S. rodhaini* clearly indicate that the East African *S. mansoni* samples from Tanzania and Uganda are sister to a clade containing all other *S. mansoni*, including those from the Caribbean, Brazil, Senegal, Niger, and Cameroon. Our data supports previous work from whole genomes, mitochondrial genes, and microsatellites suggesting that *S. mansoni* emerged in East Africa (Crellen et al., 2016; Morgan et al., 2005; Webster et al., 2013). In addition, our estimates of *Ne* are in within a range book-ended by other whole genome studies (Berger et al., 2021; Crellen et al., 2016).

The rapid decay in LD observed in E. Africa compared with W. African and American populations provides further evidence that East African *S. mansoni* populations are ancestral. Similar reductions in rate of LD decay have been observed in humans and malaria parasites, outside of their ancestral Africa range (Anderson et al., 2000; Gurdasani et al., 2015; Neafsey et al., 2008). Rapid breakdown in LD also has important practical applications for genome wide association analyses, because it allows mapping of phenotypic traits to very narrow regions of the genome (Mackay & Huang, 2018). There are multiple biomedically important traits of interest that vary in *S. mansoni* populations, including drug susceptibility or resistance, host-specificity and cercarial production (Anderson, LoVerde, Le Clec’h, & Chevalier, 2018). We recently used GWAS for mapping resistance to the first line drug (Praziquantel) in laboratory schistosome populations (Le Clec’h et al., 2021). Most laboratory *S. mansoni* populations tested were from S. America, where LD decays relatively slowly: these are not ideal for GWAS. Establishment of laboratory *S. mansoni* populations from East Africa, or GWAS analyses using parasites directly from the field would be valuable for future GWAS with *S. mansoni*.

There is minimal allele sharing or migration between East African and other *S. mansoni* populations. F_ST_ comparisons that include Tanzania are greater than other comparisons, (with Tanzania - F_ST_ = 0.355; excluding Tanzania - F_ST_ = 0.206). East African populations are among the most strongly differentiated populations in the PCA analyses (Figure 4B) and the East African population component in ADMIXTURE analyses is absent, or at minimal levels, in other *S. mansoni* populations. Finally, mitochondrial haplotypes in Uganda and Tanzania form a distinct haplogroup from other *S. mansoni* populations. These data indicate that, while E. Africa is the likely origin of *S. mansoni*, migration or allele sharing between E. Africa and other populations is restricted.

We do not expect that human movement is a major barrier between East and West African schistosome populations. However, differences in snail – schistosome compatibility in East and West Africa may provide barriers to gene flow. Sympatric host–parasite combinations tend to show greater compatibility than allopatric combinations. This is seen in multiple host parasite systems including *Daphnia*–microsporidia (Ebert, 1994), trematode infections of snails (Lively, 1989) and minnows (Ballabeni & Ward, 1993). Strong host-specificity exists within the *Biomphalaria* and *S. mansoni* system (Mitta et al., 2017; Theron et al., 2014; Webster & Woolhouse, 1998) and a review shows that compatibility is greater between sympatric *Biomphalaria*-*S. mansoni* combinations (Morand, Manning, & Woolhouse, 1996). Sympatric schistosome–snail combinations result in rapid immune suppression and rapid parasite development, while allopatric schistosome-snail combinations result in a slower immune cell proliferation and a non-specific generalized immune response which reduced parasite growth and establishment (Portet et al., 2019). There is a developing understanding of *Biomphalaria* phylogenetics (Jorgenson, Kristensen, & Stothard, 2007), phylogeography (Dejong et al., 2003), and compatibility relationships among East African snail species (*B sudanica*, *B. pffeiferi* and *B. choanomphala*) and *S. mansoni* (Mutuku et al., 2021; Mutuku et al., 2017), but further research is needed to understand compatibility of allopatric snail–schistosome combinations from East and West Africa. We suggest that the presence of fine-scale geographic structure of *Biomphalaria* populations (Webster et al., 2001) and local adaptation in sympatric *Biompahalaria*-schistosome combinations may limit parasite geneflow between E. African and W. Africa.

PCA analyses differentiate *S. mansoni* populations on an East-to-West gradient along PC2 (Figure 4B). Mitochondrial haplotypes in Senegal and Cameroon are intermediate to Tanzania and Senegal (Figure 6A). The species tree from autosomal SNV data (Figure 6A) indicates that *S. mansoni* is in a series of nested, well-supported clades from East Africa (Tanzania + Uganda) to Cameroon to West Africa (Niger + Senegal). F_ST_ and *r* (Mantel) values between Tanzania, Niger, and Senegal reflect increasing isolation with distance across Africa. These observations combined with the origination of *S. mansoni* in East Africa confirms an East-to-West, stepwise expansion of *S. mansoni* from Tanzania and Uganda → Cameroon → Niger → Senegal (Crellen et al., 2016; Morgan et al., 2005; Webster et al., 2013).

*Does Hybridization between* S. rodhaini *and* S. mansoni *contribute to elevated East African Diversity* – Several closely related *Schistosoma* species are able to hybridize with the production of viable offspring confirmed via experimental rodent infections. The potential for hybridization between animal and human *Schistosoma* species is a significant public health concern (Borlase et al., 2021; Léger et al., 2020; Leger & Webster, 2017; Stothard, Kayuni, Al-Harbi, Musaya, & Webster, 2020). Our group, and others, have recently shown that ancient hybridization and adaptive introgression has resulted in the transfer of genes from the livestock species *Schistosoma bovis* into *S. haematobium*: west African *S. haematobium* genomes contain 3-8% introgressed *S. bovis* sequences and *S. bovis* alleles have reached high frequency in some genome regions (Platt et al., 2019; Rey, Toulza, et al., 2021). The sister species of *S. mansoni*, *S. rodhaini*, parasitizes rodents and is primarily located in Eastern Africa (Rey, Webster, et al., 2021). *S mansoni* and *S. rodhaini* have been shown to readily hybridize in and produce fertile offspring in the lab (Théron, 1989). Natural hybrids have been reported in Kenya and Tanzania (Morgan et al., 2003; M. Steinauer et al., 2008), although hybrids have only been detected from their snail intermediate host and never encountered in the mammalian hosts humans and rodents (Rey, Webster, et al., 2021). We were unable to find evidence of hybridization in 55 samples collected from Tanzania. Both species are clearly separated in genotypic space with differences between the species accounting for the largest component in the PCA (Figure 4A; PC1=34.7% variation) and F_ST_ between *S. rodhaini* and *S. mansoni* populations is very high (F_ST_ = 0.912). Admixture analyses also clearly differentiated *S. rodhaini* from all other *S. mansoni* populations (Figure 5) and we were unable to identify admixture signal between *S. rodhaini* and the Tanzanian population with genome-wide statistics including D, D_3_, or F_3_, (Table 3).

Hybridization between these two species is thought to be rare (≤7.2%) (Morgan et al., 2003; Rey, Webster, et al., 2021; M. Steinauer et al., 2008; M Steinauer et al., 2008). Our sample size may not be large enough to identify rare hybrids. Further we analyzed exome (coding) data, which may underrepresent introgressed alleles if they are selected against. These caveats aside, our analyses clearly fail to identify recent hybridization between *S. mansoni* and *S. rodhaini*. We conclude that *S. rodhaini* introgression does not contribute to the high, genetic diversity in our Tanzanian *S. mansoni* samples.

### Expansion into the Americas

Previous work has shown that *S. mansoni* was exported from Africa to the Americas during the Trans-Atlantic slave trade (Crellen et al., 2016; Desprès et al., 1993; Files, 1951; Fletcher et al., 1981; Morgan et al., 2005; Webster et al., 2013). Here, we use genomic data to investigate the likely source population(s), number of introductions, evidence for bottlenecks and parasite adaptation during colonization.

#### Source populations

Of the two West African populations sampled (Niger and Senegal), our results support stronger relationships between Brazil and Niger, than with Senegal. While the species tree (Figure 6B) appears to rule out Niger or Senegal as the direct source population for Brazilian *S. mansoni*, there is evidence of allele sharing between the Nigerian and Brazil populations. First, the dominant mtDNA haplotype in Brazilian and Nigerien samples are shared (Figure 6A). Second, every Nigerien

population contains at least 10.1% of the Brazilian component as shown in the Admixture analyses (mean 15.8%; Figure 5). Third, Niger and Brazil are more closely associated with each other in the genotypic continuum represented by PC2 than Brazil is to other African populations (Figure 4B). Finally, TreeMix identified a single weak migration edge between Brazil and Niger (Figure 6C) confirming a relationship between these two populations.

A simple hypothesis from the data is that, assuming the general East-to-West expansion holds at finer geographic scales, the source population that was eventually exported to Brazil, likely occurs somewhere between Benin and Angola. These countries fell within the Bight of Benin, Bight of Biafra, and West Central Africa slave trading regions (Figure 8). Our Brazilian samples were collected in Ponto dos Volantes in Minas Gerais, Brazil. This location is relatively equidistant from major slave ports in Bahia (527 Km) and around Rio de Janeiro (706 Km). In all, more than 3.5 million (M) enslaved peoples were imported into Brazil (Supplemental Table 7; Slave Voyages Database, 2009). Of all the enslaved people exported to Brazil, 82% disembarked at ports in either Bahia (1.3M) or Rio de Janerio (1.5M). In Africa, slave exporting markets in the Bights of Benin and Biafra and West Central Africa were responsible for 53.6%, 5.0% and 33.8% of peoples exported to Bahia and 1.3%, 1.1% and 74.9% of peoples exported to south east Brazilian ports (Slave Voyages Database, 2009). Taken together these data imply that the Brazilian population we sampled in Ponto dos Volantes most likely originated from markets in the Bight of Benin or West Central Africa. In our phylogenetic analyses, the Brazilian population falls within a clade containing samples from Senegal, Niger, and the Caribbean but excludes the single Cameroonian sample. If the Cameroonian sample is representative of the Cameroonian population, then it may be that Cameroon and Niger represent the eastern and western limits of the unrepresented source population. This area more closely aligns with the Bight of Benin, a region containing parts of Nigeria and Benin. Additional samples from these regions are needed to test this hypothesis. It is also important to note that, the Brazilian samples here represent a single, geographic location (Ponto dos Volantes) and that the source for this population may not extrapolate to larger regions, or even to other locations in Brazil.

**Figure 8.**
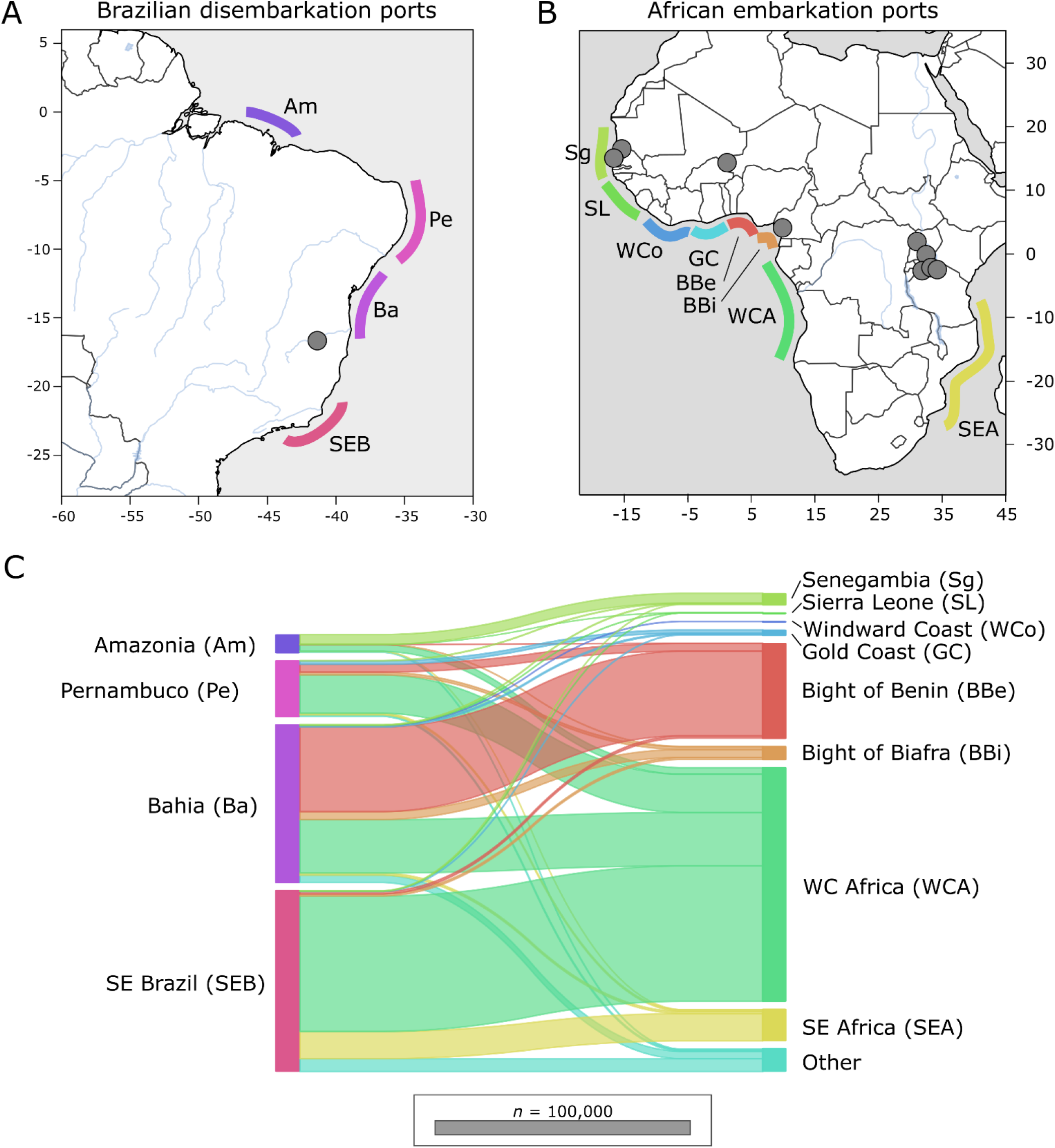
Export and importation of slaves between Africa and Brazil. During the Transatlantic Slave Trade, more than 12 million people were trafficked from Africa. Of these, 3.5 million were exported into Brazil at major ports along the east coast. Most peoples originated from regions along the West African coast from Benin to Angola. General locations of major embarkation and disembarkation ports shown along each coastline. Collection localities for *S. mansoni* are represented by grey dots. Slave exportation data is from the Voyages Database (2009).

#### No evidence for population bottlenecks during colonization

Previous mtDNA analyses have shown reduced diversity in S. American parasites (Desprès et al., 1993; Fletcher et al., 1981; Morgan et al., 2005; Webster et al., 2013) suggesting bottlenecks during colonization. Our data is consistent with this showing a 2-3-fold reduction in mtDNA in Brazil compared to W. African parasites. However, genome-wide summary statistics of autosomal sequence data tell a different story, and provide no evidence for population bottlenecks associated with *S. mansoni* introduction and establishment in Ponto dos Volantes, Brazil (Table 1). Nucleotide diversity, measured by π, was higher in Brazil than either of the West African populations (Niger or Senegal), perhaps because the Brazilian population is derived from several west African source populations. The Brazilian *N_e_* was roughly the same as Niger and comparable to Senegal. Mean Tajima’s *D* calculated from genome-wide, SNV data is negative in each of the African populations and highest in the Caribbean samples (mean Tajima’s *D* = 0.929; Supplemental Table 2). Tajima’s *D* values in our study using exome SNV data from different regions of Niger, Senegal and Tanzania were comparable with values calculated with whole genome SNV data from Ugandan *S. mansoni* (Berger et al., 2021). By contrast mean Tajima’s *D* in Brazilian samples is close to 0 (mean Tajima’s *D* = 0.034; 95% CI [-0.18 to -.085]) and is significantly greater than mean Tajima’s *D* in the African samples (T-Test *p*<0.001; Supplemental Table 2).

The nuclear genomic data clearly suggest that the establishment of *S. mansoni* in Brazil was not associated with a significant population bottleneck and had minimal impacts on genome-wide levels of genetic diversity. The discrepancy between mtDNA and nuclear DNA may stem from two sources. First, mtDNA has an effective population size ¼ that of nuclear genes (Birky, Maruyama, & Fuerst, 1983), and can potentially provide a more sensitive indicator of bottlenecks. Second, and perhaps more critical, mtDNA constitutes a single marker, so may poorly reflect population history (Anderson, 2001). Extensive laboratory passage may also result in bias in population summary statistics. For example, the Caribbean samples examined have undergone 2-15 generations of lab passage which is probably responsible for the elevated Tajima’s *D* at in this population (Crellen et al., 2016).

#### Number of introductions

All Brazilian and Caribbean samples are paraphyletic fall between the Cameroonian sample and West African clade in Figure 6B. The relationships among these samples are resolved but not supported outside of a monophyletic clade containing all Brazilian samples. As a result, the autosomal phylogeny by itself does not conclusively support one or multiple introductions into the Americas. The two Caribbean samples from Guadeloupe island contain unique mitochondrial haplotypes absent from Brazilian (Figure 6A); however, this is, at best, only weak evidence for independent introductions into Brazil and the Caribbean from the data we have available. The higher autosomal diversity of Brazilian *S. mansoni* compared with two sampled West African populations provides additional indirect evidence for multiple origins.

*Plasmodium falciparum* and *Wuchereria bancrofti* became established in the Americas at the same time as *S. mansoni*, and without apparent bottlenecks (Small et al., 2019; Yalcindag et al., 2012). In both cases it is hypothesized that high levels of diversity were maintained by recurring introductions from multiple sources (Rodrigues et al., 2018; Small et al., 2019; Yalcindag et al., 2012). More than 3.5 M enslaved peoples were exported into Brazil from across the continent of Africa (Supplemental Table 7; Slave Voyages Database, 2009). *S. mansoni* prevalence varies widely between sites in western and central Africa, with estimates ranging from 0.3% (Gambia; Sanneh et al., 2017) to 89% (Democratic Republic of the Congo; Kabongo et al., 2018). Assuming *S. mansoni* prevalence during the period of the Atlantic Slave Trade is comparable to current levels and that infected persons contained multiple *S. mansoni* individuals (genotypes; Van den Broeck et al., 2014), it is plausible that hundreds of thousands, and more likely millions, of reproductively viable *S. mansoni* were introduced into Brazil from across Africa. As a result, individual genotypes that may have been separated by thousands of miles in Africa were brought into close contact in the Americas and would lead to higher levels of diversity in there than in any single population in Africa.

#### Adaptation during colonization

We hypothesized that *S. mansoni* introduced into the Americas would have been exposed to novel selective pressures as they adapted to new biotic and abiotic challenges. For example, the *S. mansoni* life cycle requires an intermediate snail host in which miracidia mature into cercariae that are capable of infecting humans. In Africa, *S. mansoni* use *Biomphalaria alexandrina*, *camerunensis*, *choanomphala*, *pfeifferi*, *stanleyi*, and / or *sudanica* as the snail host, none of which are present in the Americas. Instead, these parasites have adapted to using different *Biomphalaria* hosts, including *B. glabrata*, *B. straminea*, and *B. tenagophila* (Figure 2; Hailegebriel, Nibret, & Munshea, 2020; Kengne-Fokam, Nana-Djeunga, Bagayan, & Njiokou, 2018; Vidigal et al., 2000). We examined exomic SNV data to identify genes and larger regions of the genome under selection at a finer scale and identified 0-5 putative regions of selection from each of the major populations (Table 4).

In the Brazilian samples, we identified five putative selected regions that contain 126 genes (Supplemental Table 6). π and Tajima’s *D* was significantly reduced in these five regions compared to genome wide averages which is expected if these loci are, or have been, under selection (Supplemental Table 3). One region is shared between the Brazilian and Senegalese populations. Forty-six genes fall within this Sengal:Brazil overlapping region, leaving 80 genes and 4 loci that are likely experiencing population specific, positive selection specific. Even within the group of 80 genes, there are 9 with strong signals of selection (Table 5). These genes contained variants with *H* and LR values in the 99^th^ percentile in addition to being greater than the threshold defined by neutral simulations. Several genes within this group are associated with housekeeping functions, including transcription and protein degradation (Smp_060090, Smp_162000, and Smp_246630). Two uncharacterized proteins were identified (Smp_341570 and Smp_123590) but we are not able to speculate on their function. Of particular interest are two possible transcription factors, an uncharacterized protein containing a *helix, loop, helix* domain (Smp_123570) and a putative TATA-box binding protein (Smp_073680). It is possible that adaptation to the Brazilian environment was driven by changes in gene expression, however more work is needed to understand the potential role of these loci in adaptation to the Americas.

### Selection on African <u>S. mansoni</u>

We examined selection on African *S. mansoni* as part of the process to identify unique signals of selection in the Brazilian population. We identified 112 and 157 genes under selection in 3 regions each for the Nigerien and Senegalese populations (Table 4). One of the three regions, and 22 genes, were shared between Niger and Senegal. We failed to identify any regions of selection in Tanzania using our combined criteria, however results from individual tests of selection (H-Scan, SweepFinder2) did overlap at nine of 25 regions identified in large Ugandan by Berger et al. (2021) (Supplemental Table 8). These regions were identified using a variety of within (iHS) and between population (FST, XP-EHH) tests on miracidia isolated from two Kenyan locations with differing histories of praziquantel treatment.

## Conclusions

Our analyses identified an East-to-West expansion of *S. mansoni* across Africa. Sometime during this expansion one or more Central African population(s), likely located between Angola and Benin, were transported to Brazil. Genome-wide signatures of diversity, measures of allele frequencies (Tajima’s *D*), and estimates of *Ne* are comparable between *S. mansoni* in Brazil, Niger, and Senegal and do not imply the presence of a bottleneck during the establishment of *S. mansoni* in Brazil. We did find five genome regions under selection in Brazil, 4 of which are population specific. In total, 80 genes fall within these regions and may be associated with *S. mansoni*’s adaptation to novel selection pressures associated with the Americas. We identified 9 genes with the strongest signals of selection that are candidates for future experimental work.

## Acknowledgments

Sandra Smith and John M. Heaner (Texas Biomedical Research Institute) provided computational research support. Matthew Berriman provided the most recent *S. mansoni* genome sequence and annotation prior to publication. This research was funded by the National Institute of Allergy and Infectious Diseases (NIAD R01 AI097576-01 and NIAD 5R21AI096277-01), Texas Biomedical Research Institute Forum (award 0467). Samples acquisition was supported by funding from the Wellcome Trust (104958/Z/14/Z), the Gates Foundation coordinated by the University of Georgia Research Foundation Inc. (RR374-053/5054146 and RR374-053/4785426), and a ZELS research grant (combined BBSRC, MRC, ESRC, NERC, DSTL, and DFID: BB/L018985/1). The authors declare no competing interests.

## Author Contributions

Study design done by F.D.C., T.J.A., and W.L.C; formal analysis done by F.D.C, R.N.P., and T.J.A.; writing, original draft, done by R.N.P. and T.J.A.; reviewing and editing, done by A.G., A.E., B.W., D.R., F.D.C ., G.O., J.P.W., M.M, P.T.L., R.R.dA., R.N.P., S.K., T.J.A., and W.L.C.; investigation was done by. F.D.C., M.M., and W.L.C.; resources were provided by: F.D.C., and T.J.A.

## Data Availability

Data used in this manuscript was previously published in (Berriman et al., 2009; Chevalier et al., 2019; Crellen et al., 2016; International Helminth Genomes Consortium, 2019; Le Clec’h et al., 2021) under multiple NCBI BioProject (PRJNA439266, PRJNA560070, PRJEB522, PRJEB526, PRJNA743359 and PRJNA773498) and NCBI Short Read Archive (ERR046038, ERR103049, ERR103050, ERR119614, ERR119615, ERX284221, ERR310938, ERR539846, ERR539847, ERR539848, and ERR9974) accessions.

## Code Availability

Scripts, notebooks, and environmental yaml files are available at https://github.com/nealplatt/sch_man_nwinvasion/releases/tag/v0.2 (last accessed 21 Oct 2021) or https://doi.org/10.5281/zenodo.5590460 (last accessed 21 Oct 2021).

