## Supplemental Figures 1-3 for "Genomic analysis of a parasite invasion: colonization of the Americas by the blood fluke, *Schistosoma mansoni*"

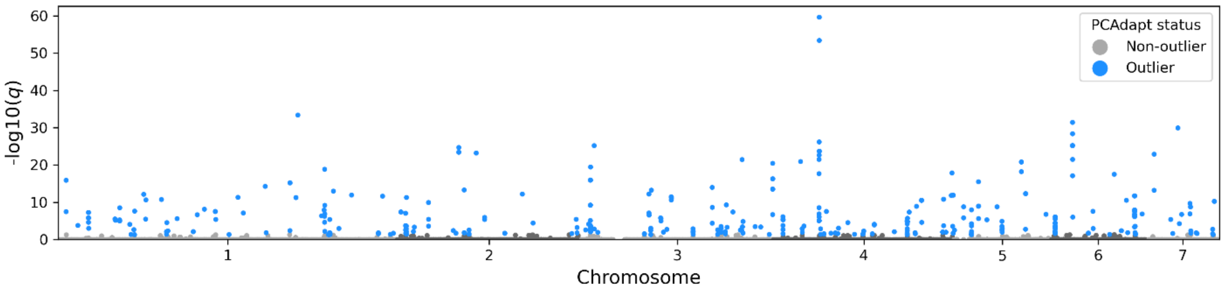


**Supplemental Figure 1. Directional selection across three *S. mansoni* populations** – Population specific, directional selection was estimated at each single nucleotide variant (SNV) across the exome using pcadapt v4.3.3 (Luu et al., 2017). We examined three *S. mansoni* populations including indiviudals from Niger, Senegal, and Brazil. Outlier SNVs were identified after multiple test correction (Bonferroni) and α = 0.05.

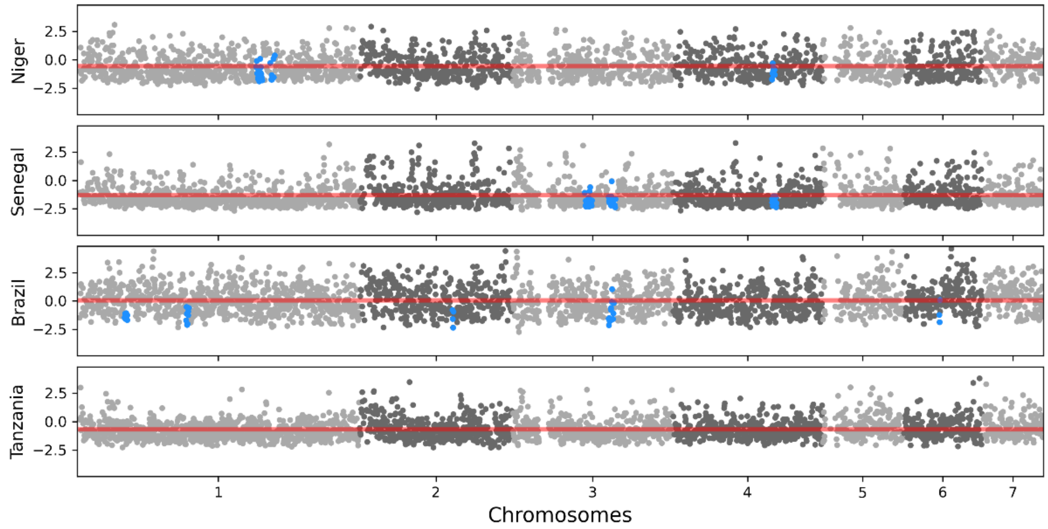


**Supplemental Figure 2. Nucleotide diversity (π) across the genome.** π was measured across the genome in each *S. mansoni* population. Average π for each population is indicated by the red line. π in regions identified as putative regions of selection are shown in blue. π was significantly lower in putative regions of selection than expected based on genome-wide averages in Niger, Senegal, and Brazil. We did not identify any putative regions of selection in the Tanzanian population.

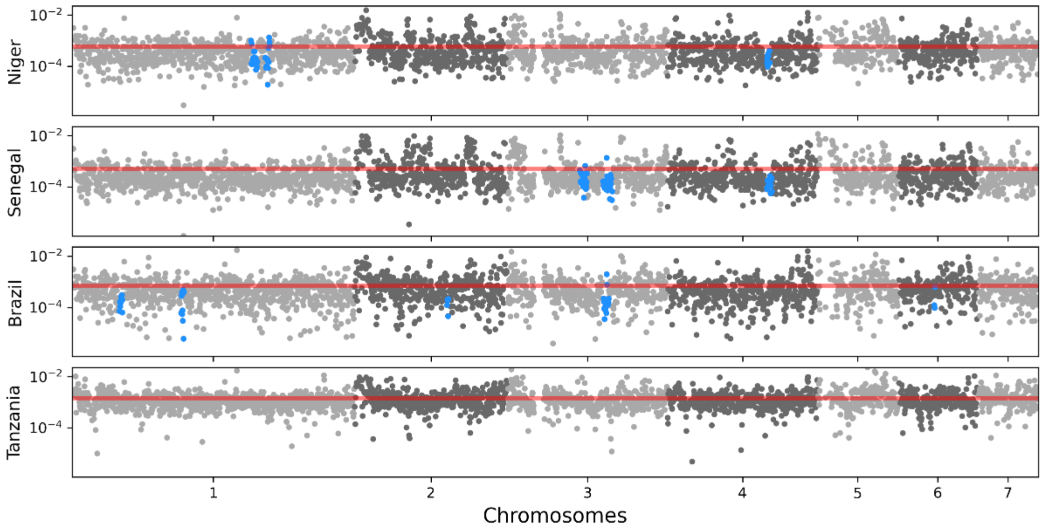


**Supplemental Figure 3. Tajma's *D* across the genome.** π was measured across the genome in each *S. mansoni* population. Average Tajma's *D* for each population is indicated by the red line. Tajma's *D* in regions identified as putative regions of selection are shown in blue. Tajma's *D* was significantly lower in putative regions of selection than expected based on genome-wide averages in Niger, Senegal, and Brazil. We did not identify any putative regions of selection in the Tanzanian population.
